# In vitro modelling of anterior primitive streak patterning with human pluripotent stem cells identifies the path to notochord progenitors

**DOI:** 10.1101/2023.06.01.543323

**Authors:** M. Robles-Garcia, C. Thimonier, K. Angoura, E. Ozga, H. MacPherson, G. Blin

**Affiliations:** Institute for Regeneration and Repair, Institute for Stem Cell Research, School of Biological Sciences, The University of Edinburgh, 5 Little France Drive, Edinburgh BioQuarter, Edinburgh EH16 4UU, UK

## Abstract

Notochord progenitors (NotoPs) represent a scarce yet crucial embryonic cell population, playing important roles in embryo patterning and eventually giving rise to the cells that form and maintain intervertebral discs. The mechanisms regulating NotoPs emergence are unclear. This knowledge gap persists due to the inherent complexity of cell fate patterning during gastrulation, particularly within the anterior primitive streak (APS), where NotoPs first arise alongside other important progenitors including neuro-mesodermal and endodermal progenitors.

To gain insights into this process, we use micropatterning together with FGF and the WNT pathway activator CHIR9901, to guide the development of human embryonic stem cells into reproducible patterns of APS cell fates. We show that small variations in CHIR9901 dosage dictate the downstream dynamics of endogenous TGFbeta signalling which in turn controls cell fate decisions. We show that sustained NODAL signalling induces endoderm while NODAL inhibition is needed for NMP specification. Furthermore, we unveil a crosstalk between TGFbeta and WNT signaling pathways, wherein TGFbeta inhibition enhances WNT activity. Finally, we demonstrate that the timely inhibition of TGFbeta signalling is imperative for the emergence of NotoPs.

Our work elucidates the signalling regimes underpinning NotoPs emergence and provides novel insights into the regulatory mechanisms controlling the balance of APS cell fates during gastrulation.

## Introduction

In vertebrate embryos, the tissues of the posterior axis, including the spinal cord, the cartilage, bones and muscles of the spine, as well as the gut, are all laid down progressively in an anterior-to-posterior direction. This evolutionary-conserved process, termed axial elongation (reviewed in (Henrique et al., 2015; Neijts et al., 2014; Wymeersch et al., 2021), is governed by lineage-restricted progenitors emerging during gastrulation in the anterior portion of the primitive streak (APS) (Fig1A).

**Figure 1.**
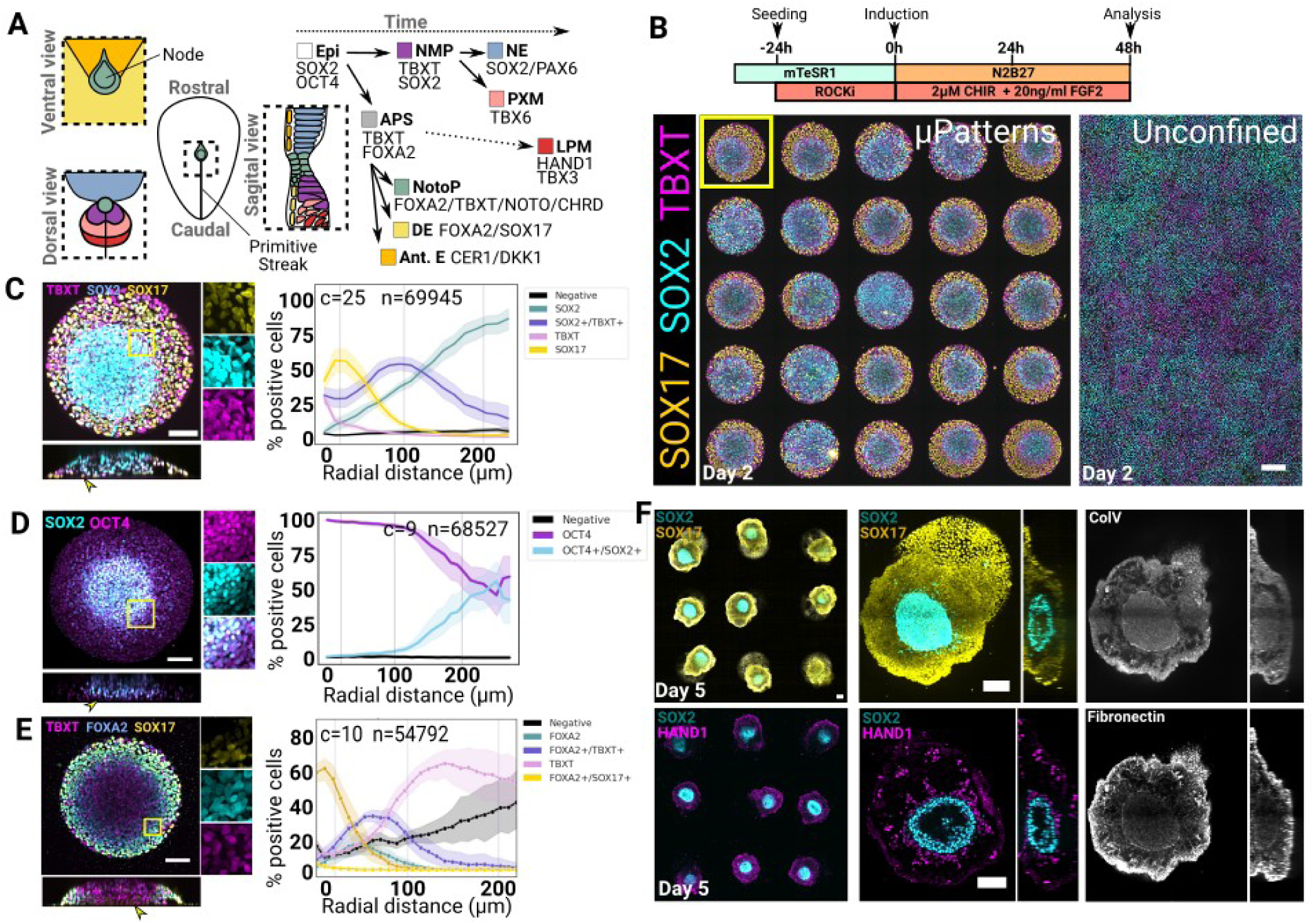
Geometrical confinement elicits radial patterning of endoderm and mesoderm biased cell fates. **A** Diagrammatic representation of the putative organisation and hierarchies of cell fates around the node in a human embryo at the end of gastrulation (left). Extra-embryonic structures are omitted. Relevant cell lineages are indicated together with the associated cell fate markers used in this study (Right). **B** Illustration of the protocol used in C to E (Top) and confocal max projections showing unconstrained and micropatterned cultures (Bottom). The yellow outline indicate the colony shown at higher resolution in C. Scale bar: 200µm. **C-E** Confocal max projections of micropatterned colonies stained at 48h, scale bar: 100µm. The corresponding Z-projection is shown below each image with yellow arrowheads pointing at SOX17+ or SOX2-cells lining the central domain at the bottom of the colony. Radial profiles of cell type abundance are shown on the right. c: number of analysed colonies, n: number of analysed nuclei, shaded area: 95% confidence interval. **F** Confocal images of colonies stained at day 5 of differentiation. Wide fields are shown on the left and a representative colony on the right. Images are max projections except for the selected colony stained with HAND1, ColV and Fibronectin where a z-slice was selected to better show the localisation of the positive signal. Scale bar: 100µm. APS: anterior primitive streak, Epi: Epiblast, NotoP: Notochord Progenitors, NMPs: Neuromesodermal progenitors. NE: Neurectoderm, PXM: paraxial mesoderm, LPM: Lateral plate mesoderm, Ant.E: Anterior endoderm.

Among these, notochord progenitors (NotoPs) give rise to the notochord, a cylindrical structure that forms along the ventral midline in chordate embryos (Balmer et al., 2016; Stemple, 2005). Work in animal models has shown that the notochord ensures essential functions during development, both as a source of localised signalling molecules necessary for the proper patterning of adjacent tissues (Streit and Stern, 1999) and as a mechanical structure contributing to the straightness of the rostro-caudal axis as the embryo elongates (Bagnat and Gray, 2020; McLaren and Steventon, 2021). Eventually, the cells that form the notochord contribute to the nucleus pulposus (Choi et al., 2008; McCann et al., 2012), the central region of intervertebral discs responsible for the homeostasis of the surrounding fibro-cartilagenous tissue (reviewed in (Wise et al., 2020)).

Previous work in mouse embryos has shown that the specification of the notochord requires the cooperation of WNT and NODAL signalling (Dunn et al., 2004; Lickert et al., 2002; Merrill et al., 2004; Vincent et al., 2003; Yamamoto et al., 2001). However, the exact signalling requirements for the specification of NotoPs has remained elusive. Indeed, in spite of previous efforts (Colombier et al., 2020; Diaz-Hernandez et al., 2020; Winzi et al., 2011; Zhang et al., 2020), the derivation of NotoPs from pluripotent stem cells remains inefficient unless transcription factors regulating the notochord fate are overexpressed and even these conditions lead to mixed population of cells including alternative cell fates from the mesoderm and endoderm lineage (Schifferl et al., 2023; Warin et al., 2024). Furthermore, apart from one exception (Xu et al., 2021), 3D organoids that mimic axial elongation also lack notochord in spite of the fact that WNT and NODAL signalling are both active in these in vitro models (Beccari et al., 2018; Cermola et al., 2021; Moris et al., 2020; Turner et al., 2017) suggesting there may be additional unknown cues needed for notochord specification.

This gap in understanding stems from the relative rarity of NotoPs in the embryo and from the complexity and rapidity of the cell fate decisions during gastrulation. In mouse embryos, NotoPs emerge in close proximity to both the definitive endoderm (Beddington and Robertson, 1989; Scheibner et al., 2021), and neuromesodermal progenitors (NMPs), the bi-fated cells that form the somitic mesoderm and the spinal cord (Cambray and Wilson, 2002; Cambray and Wilson, 2007; Forlani et al., 2003; Koch et al., 2017; Tzouanacou et al., 2009). All three populations, endoderm, NotoPs and NMPs, have been reported to emerge at around mid-gastrulation (Ang and Rossant, 1994; Cambray and Wilson, 2002; Pour et al., 2022; Scheibner et al., 2021; Sulik et al., 1994; Yamanaka et al., 2007), eventually forming a progenitor zone organised along the antero-posterior axis of the embryo. On the anterior side of the primitive streak, endodermal progenitors undergo partial epithelial to mesenchymal transition and intercalate into the underlying visceral endoderm to form the gut endoderm (Kwon et al., 2008; Scheibner et al., 2021; Viotti et al., 2014). NotoPs establish the ventral epithelial layer of the node, demarcating the rostral and caudal regions of the embryo (Bakker et al., 2016; Kinder et al., 2001). NMPs populate the epithelial space directly adjacent and posterior to the node and together with NotoPs form the progenitor growth zone that fuels axial elongation (Fig 1A) (Abdelkhalek et al., 2004; Wymeersch et al., 2019). How the patterning and balanced proportion of these populations is established is not understood.

Here, we set out to identify the signalling requirements distinguishing NotoPs from other APS cell fates in a human context. To tackle this challenge and circumvent the technical and ethical limitations inherent to research on rare embryonic cell populations, we use micropatterning to guide the development of human embryonic stem cells (hESC) into reproducible patterns of APS cell fates. We start with an NMP-inducing medium containing FGF and CHIR99021 (CHIR), a potent activator of the WNT pathway (Cohen and Goedert, 2004). First, we show that in contrast to cells grown as a 2D monolayer culture, cells grown on micropatterns preferentially differentiate towards mes-endodermal lineages. We show that partial inhibition of endogenous NODAL signalling prevents endoderm specification and restores the emergence of NMPs but fails to induce NotoPs. Then we uncover that hESC grown on micropatterns respond non-monotonically to increasing doses of CHIR with distinct downstream temporal dynamics of NODAL signalling. We also identify a crosstalk between WNT and NODAL wherein NODAL inhibition potentiates WNT signalling activity. By inhibiting NODAL and BMP signalling 24h post CHIR induction when the cells are still uncommitted to endoderm, we are able to efficiently redirect the cells from the endoderm to the NotoP fate.

Overall, our study uncovers signalling cross-talks and dynamics that correlate with key lineage restrictions prior to axial elongation and identifies the path to the notochord lineage.

## Results

### hESC colony confinement directs patterning of mesendoderm-biased cell fates

In order to study the mechanisms underlying NotoPs specification in a human context, we set out to establish an *in vitro* system that would mimic aspects of the formation of the axial progenitor zone using hESC. NotoPs emerge in close proximity to NMPs within a small, confined region of the embryo (Cambray and Wilson, 2007). Furthermore, scRNAseq analysis have shown that *in vitro* derived NMPs may contain a rare population of cells with a node-like signature (Edri et al., 2019). Therefore, we hypothesised that combining an NMP derivation medium (Frith and Tsakiridis, 2019; Gouti et al., 2014) with geometrical confinement (Blin, 2021) would elicit the self-organisation of hESC colonies into distinct domains of APS cell fates including NotoPs (Fig 1B)

To test this idea, we stained the cells 48h post CHIR and FGF induction for the NMP markers TBXT and SOX2 and included the endodermal marker SOX17 (Kanai-Azuma et al., 2002; Viotti et al., 2014) as a proxy for the emergence of additional APS cell fates that do not normally appear in NMP differentiation monolayers (Frith and Tsakiridis, 2019) (Fig 1B). In line with previous reports (Frith et al., 2018; Gouti et al., 2014), cells cultured in conventional 2D dishes co-expressed TBXT and SOX2 (Frith et al., 2018; Gouti et al., 2014) and remained negative for SOX17 (Fig 1B). In sharp contrast, cells grown on micropatterns consistently formed radially organised domains of cell fate markers, with a TBXT/SOX2 domain located between a SOX2-only domain in the centre and a SOX17 domain at the periphery (Fig 1B and 1C). We confirmed that this phenomenon was reproducible across several human pluripotent cell lines (Sup Fig 1).

We next tested whether different colony sizes may change the proportion of cell fates on micropatterns (Sup Fig 2A). (Kwon et al., 2008; Scheibner et al., 2021; Viotti et al., 2014)Using our established quantitative immunofluorescence pipeline (Blin et al., 2019; Wisniewski et al., 2019), we measured the proportion of each individual population (Sup Fig 2B) and plotted these proportions as a function of the radial distance from the colony edge (Sup Fig 2C). We found that all three domains, i.e SOX17, TBXT/SOX2 and SOX2-only domains, remained located at a consistent distance from the colony edge across all colony diameters except for colonies below 320µm where this rule did not apply as strictly. Incidently, the central domain of SOX2 expressing cells increased in size proportionally with colony diameter effectively increasing the percentage of SOX2-only cells. These observations may suggest that the mechanisms driving the radial cell fate organisation in this context is boundary-driven as reported in other micropatterned colony systems (Etoc et al., 2016; Martyn et al., 2019; Warmflash et al., 2014). For the subsequent experiments, we decided to use 500µm colonies because this diameter offered a good compromise for analysis and imaging with a clear radial fate marker distribution (Fig 1 C).

**Fig 2.**
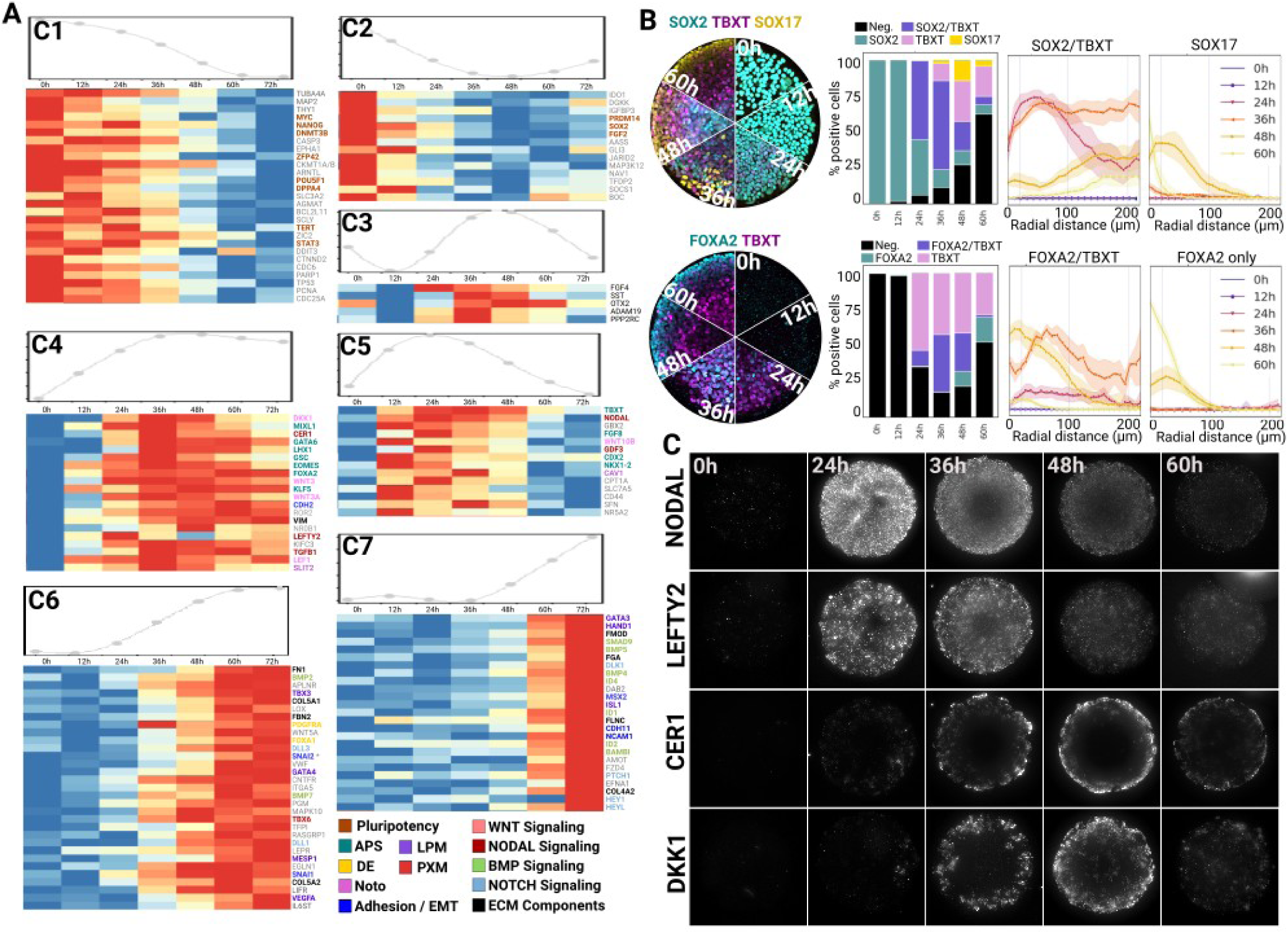
Dynamics of NODAL, WNT and BMP signalling correlate with the loss of axial cell fates and the emergence of definitive endoderm and lateral plate mesoderm. **A** Nanostring time course analysis. Cells were treated as in Fig. 1B. For each cluster, a model spline of the cluster is shown at the top of the gene expression heatmap. Genes are ordered from top to bottom in decreasing order of maximum differential expression. Colours represent the mRNA count normalised on a per gene basis. B Time course analysis via quantitative immunofluorescence of 500µm colonies. Left: montage of confocal max projections, Middle: stacked bar plot showing the relative proportions of individual cell populations over time, Right: Radial profiles of the percentage of cells expressing the markers indicated at the top of the plots. C Widefield images of 500µm colonies stained via branched DNA FISH at selected time points.

We next asked about the developmental state of the cells forming the central SOX2 domain. In mouse embryos, SOX2 is initially co-expressed with OCT4 in the pluripotent epiblast and remains expressed in the developing neurectoderm while OCT4 becomes progressively lost as the cells exit pluripotency (Avilion et al., 2003; Osorno et al., 2012). In our colonies, OCT4 was still expressed in the central SOX2 domain at this stage indicating that the cells had not yet exited pluripotency (Fig 1D).

Next, we tested for the presence of NotoPs. We first looked for the co-expression of FOXA2 and TBXT which are both essential for the development of the notochord (Ang and Rossant, 1994; Lolas et al., 2014; Tamplin et al., 2011; Yamanaka et al., 2007). Cells expressing FOXA2 could be found all around the colony spanning a domain of approximately 100µm from the edge (Fig 1E). This domain could be divided into a SOX17+ outter domain indicative of the endodermal fate and a FOXA2/TBXT inner domain co-localising with the SOX2/TBXT domain shown in Fig 1C. Since these markers are also transiently co-expressed in the nascent mesendoderm during gastrulation (Burtscher and Lickert, 2009), we performed FISH against the NotoP-specific marker NOTO (Abdelkhalek et al., 2004; Plouhinec et al., 2004). We did not observe any positive cells for this marker suggesting that the TBXT/FOXA2+ cells in these colonies likely represent an early mesendoderm population at 48h.

To further characterise lineages emerging in micropatterned colonies, we repeated the same protocol as in Fig 1B and cultured the cells for an additional 3 days in unsupplemented N2B27 (Fig 1F). Over time, the colonies established a 3-dimensional structure composed of a cavity-comprising SOX2-positive core surrounded by a mass of SOX17 endodermal cells and clusters of HAND1+ cells. The existence of HAND1+ cells in these colonies indicate that some of the cells differentiated either to lateral plate mesoderm or extraembryonic mesoderm (Pham et al., 2022). On the other hand, we were unable to find evidence of notochord-like cells co-expressing TBXT, FOXA2 and SOX9 which is normally found in the notochord (Bagheri-Fam et al., 2006), confirming that the TBXT/FOXA2+ population identified at 48h failed to engage in the notochord lineage. We were also unable to find TBXT/SOX2 double positive cells in these structures suggesting that NMPs were also absent.

The 3-dimensional multi-tissue architecture observed here was remarkably consistent across colonies (Fig 1F). Interestingly, colonies adopted a 3-dimensional organisation as early as 48h with the formation of a dome-like structure where SOX2+ cells were elevated in comparison to other cell types (Fig 1C, D, E and Sup Fig2A). In particular, we consistently observed that SOX17+ cells, while abundant at the periphery, also formed a sparse epithelial layer lining the bottom of the colony (arrow heads in Fig 1C, E and Sup Fig2A), perhaps reflecting the behaviour of the nascent endoderm in vivo which undergoes partial EMT as it segregates from the mesoderm to form the gut endoderm epithelium during gastrulation (Kwon et al., 2008; Scheibner et al., 2021; Viotti et al., 2014) . Staining for Collagen V and Fibronectin at day 5 (Fig 1F), two extracellular matrix proteins expressed in the endoderm and mesoderm in human embryos (Zhao et al., 2024), revealed that the cells secreted their own extra-cellular matrix (ECM). Both proteins formed a basal membrane surrounding the SOX2+ domain while Fibronectin also formed a complex network inside the endodermal domain. These observations indicate that the cells organised their own ECM which likely contributed to maintaining the separation between cell fate domains and the overall morphogenetic process observed in the colonies.

Altogether, these initial experiments allowed us to establish an *in vitro* system where hESC organise into stereotypic patterns of cell fates including endoderm and mesoderm and undergo reproducible complex morphogenesis over time. While this system provides a good starting point, we were unable to detect NotoPs in these conditions, raising the question of which additional signals might be required to generate this cell type. Furthermore, NMPs did not persist on micropatterns despite the use of an NMP inducing medium suggesting that confinement potentiates endogenous cues that deflect the cells from axial cell fates towards alternative lineages.

### 2 Dynamics of NODAL and WNT signalling correlate with the loss of axial cell fates and the emergence of definitive endoderm and lateral plate mesoderm

To gain insights into the mechanisms driving cell fate diversification and the loss of axial lineages on micropatterns, we performed time course experiments and bulk RNA Nanostring analysis. We used a panel of probes consisting of the 780 genes included in the standard human embryonic stem cell gene panel together with 30 additional custom probes (listed in Sup Table 1). This panel covered a wide array of genes involved in differentiation, metabolism, signalling pathways and the cell cycle. In order to analyse our dataset, we used the Bioconductor package moanin (Varoquaux and Purdom, 2020) which allowed us to group individual genes based on their temporal profile (see Methods). We identified 7 clusters that are shown in Fig 2A. Cluster 1 and 2 identify genes that are expressed at the start and then progressively downregulated. As expected, these include pluripotency markers such as MYC, OCT4, DPPA4, DNMT3B and ZFP42. We found NANOG to be highest at around 36h and then lost rapidly consistently with the fact that NANOG is re-expressed in the posterior epiblast at the onset of gastrulation in mouse embryos (C et al., 2018; Hart et al., 2004; Osorno et al., 2012). We also found SOX2 in cluster 2 as a gene that is rapidly downregulated and starts to re-emerge at around 72h most likely as a result of its expression in the central domain undergoing neural differentiation (Fig 1H).

Cluster 4 and 5 identified genes that become expressed as early as 12h post-induction. Cluster 4 regrouped genes with a sustained expression after 36h while the genes in cluster 5 peaked at 36h and decreased thereafter. These two clusters revealed a clear APS signature with the expression of genes associated with the APS in mice including MIXL1 (Hart et al., 2002), LHX1 (Costello et al., 2015), EOMES (Costello et al., 2011), GSC, CDX2, FOXA2 (Burtscher and Lickert, 2009), KLF5 (Aksoy et al., 2014) as well as the NMP-associated gene NKX1-2 (Albors et al., 2018). Encouragingly we also found a peak of expression of SLIT2 and CAV1 at 36h, recently reported as markers of human embryonic notochord (Paillat et al., 2023; Warin et al., 2024). However, NMPs and NotoPs associated genes decreased over time indicating that axial progenitors failed to emerge. We confirmed this using quantitative immunofluorescence and observed that while the majority of the cells expressed TBXT/SOX2 at 36h initiating from the periphery at 24h, these markers were progressively lost in favour of SOX17 (Fig 2B and Sup Fig 3). Similarly, TBXT/FOXA2 co-expressing cells increased in proportion until 36h and were replaced by FOXA2 only cells at the periphery suggesting that some of these cells were on their way to form endoderm (Fig 2B and Sup Fig 3).

In fact, many of the genes found in cluster 4 and 5 of our Nanostring dataset are known regulators of the endodermal lineage. For example, both MIXL1 and KLF5 are required for specification of the definitive endoderm in the mouse (Aksoy et al., 2014; Hart et al., 2002; Moore-Scott et al., 2007) and LHX1, while expressed in the node, works together with OTX2 (found expressed transiently in cluster 3) to define anterior endoderm (Costello et al., 2015). Furthermore, looking at cluster 6 and 7, where genes become upregulated from 48h onwards, we could confirm the emergence of additional endodermal markers such as FOXA1 (Ang and Rossant, 1994), PDGFRA and GATA4 as well as several ECM components, some of which likely secreted by the endoderm including FN1, COL4A2, COL5A1, COL5A2, FBN2 and FLNC.

Endoderm was not the only lineage emerging in our colonies as we could observe clear evidence of mesoderm specification. GATA6 (Morrisey et al., 1996; Zhao et al., 2005) and EOMES (Costello et al., 2011), found in cluster 4, are both involved in the specification of the endoderm and the cardiac mesoderm lineage in the streak, and cluster 6 and 7 regrouped many genes associated with the lateral plate mesoderm including GATA3, HAND1, ISL1, TBX3 (Washkowitz et al., 2012) and MESP1 alongside the EMT markers SLUG, SNAIL and MSX2.

Overall, these data confirm our previous observations and show that cells on micropatterns follow a coherent developmental program. However, while the cells initially follow the route towards axial cell fates (i.e express APS markers), the cells eventually differentiate towards alternative lineages including endoderm and lateral plate mesoderm.

We thus turned our focus towards endogenous signalling pathways that may explain the observed endoderm and mesoderm differentiation. NODAL is a known driver of mesodermal and endodermal specification (Robertson, 2014) and its expression is positively regulated by the canonical WNT pathway (Ben-Haim et al., 2006; Norris et al., 2002); a pathway that we stimulate with CHIR in our cultures. Our Nanostring data showed upregulation and sustained expression of the WNT target gene LEF1 (Cadigan and Waterman, 2012) as early as 12h while NODAL expression peaked at 24h before decreasing progressively (FIG 2A). This peak of NODAL was followed by a peak of CER1 expression at 36h in line with previous report showing that NODAL induces its own inhibitor (Colombier et al., 2020; Funa et al., 2015). FISH confirmed the temporal expression profile of NODAL and CER1 (Fig2 C) and showed that NODAL expression was widespread across the colony at 24h alongside its antagonist LEFTY2. Interestingly, CER1 (Belo et al., 1997; Perea-Gomez et al., 2002) and the WNT antagonist DKK1, both expressed in the hypoblast of primate embryos (Bergmann et al., 2022) were expressed in a domain that overlapped spatially and temporally with the SOX17 domain (Fig2 A and B) indicating that at least a fraction of the endoderm emerging in micropattern colonies may adopt an hypoblast identity.

Altogether, our data revealed the implementation of a regulatory network of signalling molecules in micropatterns involving WNT, NODAL and their respective inhibitors.

### 3 Nodal inhibition rescues NMP emergence

Our results raise the possibility that endogenous NODAL signalling is responsible for the loss of axial cell fates and the emergence of definitive endoderm and lateral plate mesoderm on micropatterns. To confirm this hypothesis, we next decided to titrate NODAL activity using increasing doses of the NODAL receptors inhibitor SB431542 (SB) (Inman et al., 2002) (Fig 3A). We observed that even low amounts of SB was sufficient to strongly reduce the proportion of SOX17 and FOXA2 positive cells confirming that endogenous NODAL signalling activity is indeed responsible for the emergence of endoderm in this system (Fig 3A-1µM). Further increasing the dose of SB progressively expanded the central domain of SOX2 single positive cells restricting TBXT expression to the periphery. We also noted that higher doses of SB, depleted the FOXA2/TBXT population while maintaining a high percentage of SOX2/TBXT population at the periphery (Fig3 A 3µM and 10µM). We hypothesised that this SOX2/TBXT positive and FOXA2 negative cell population may be engaging in the NMP cell fate and could drive elongation.

**Figure 3.**
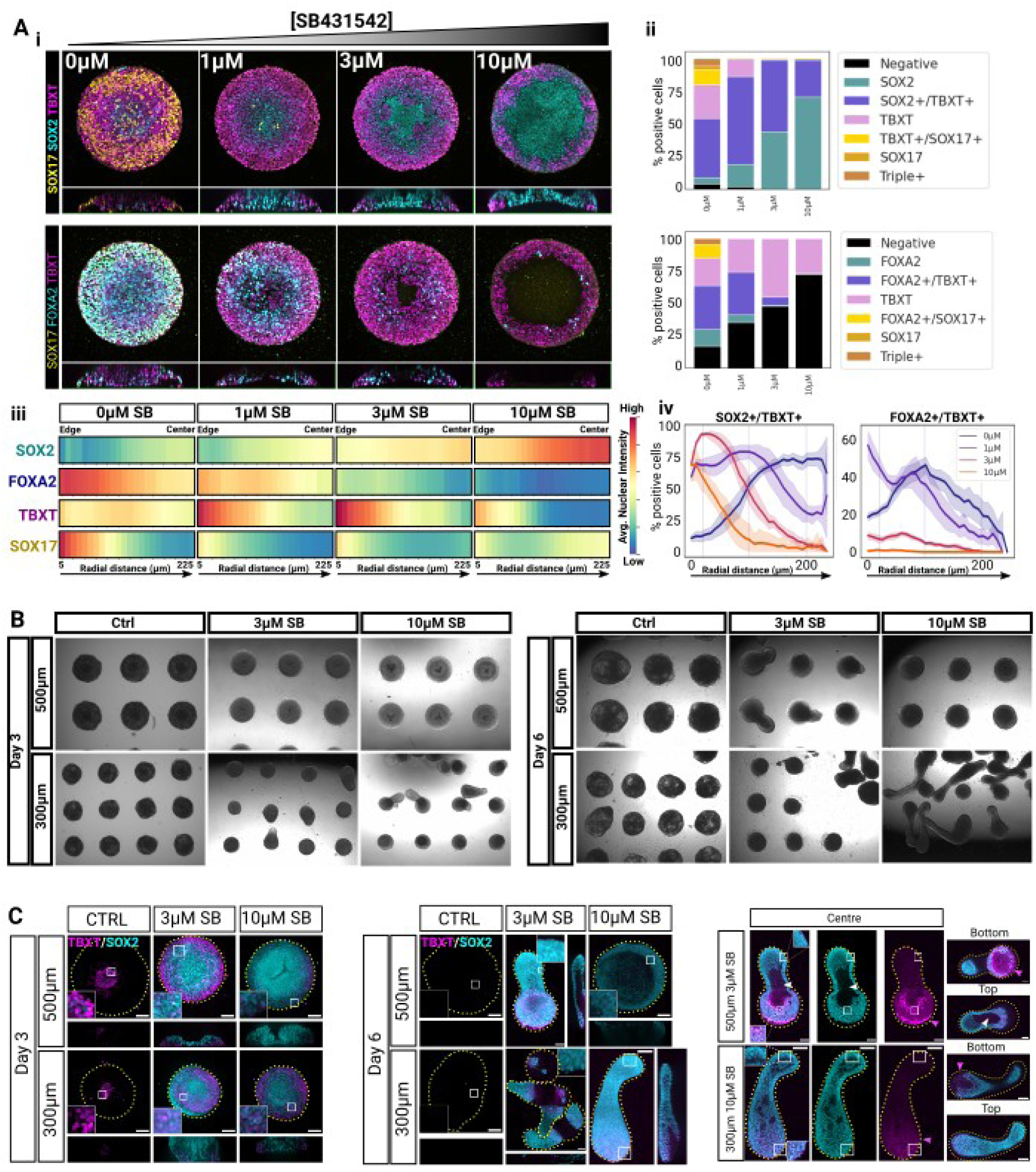
Nodal inhibition rescues NMP specification and enables tissue elongation. **A** SB dose response in 500µm colonies fixed at 48h. (i) Max projections of confocal z-stacks (top) and z-projection (bottom). (ii) Stacked bar plots showing the relative proportions of individual cell populations. (iii) Heatmaps showing the radial profile of average nuclear marker intensity scaled between 0 and 1 on a per-marker basis. (iv) Right: Radial profiles of the percentage of cells expressing the markers indicated at the top of the plots. **B** Brightfield images of hESC colonies grown on 500µm or 300µm micropatterns treated with CHIR and FGF with varying doses of the NODAL inhibitor SB431542. **C** Max projections of confocal z-stacks showing representative micropatterned colonies stained for TBXT and SOX2 (left and middle panels). The panel on the right shows selected single confocal planes of elongated structures to better show the cavity (white arrohead) and the location of the TBXT domain (magenta arrowhead). Scale bars: 100µm

To test this idea, we cultured the cells for 3 days in CHIR/FGF with or without 3µM or 10µM SB and for an additional 3 days without CHIR/FGF while maintaining SB concentration (Fig 3B and C). Since previous work with 3D models of axial elongation showed that the starting number of cells is critical to the competence of cell aggregates to elongate (Beccari et al., 2018; Martins et al., 2020; Moris et al., 2020; Olmsted and Paluh, 2021; Sanaki-Matsumiya et al., 2022; Turner et al., 2017; Veenvliet et al., 2020), we tested two different colony diameters (500µm and 300µm). As expected, we did not observe elongation in the control with either of these diameters but we did see elongation in SB containing conditions. To our surprise, we observed that competence to elongate was a function of both colony diameter and SB concentration, with elongation occurring consistently across colonies only with 3µM SB on 500µM micropatterns and with 10µM SB on 300µm micropatterns (Fig 3B and C). We next checked whether SOX2/TBXT double positive putative NMPs were maintained in these structures (Fig 3C). Indeed, we observed that elongation correlated with the conditions favouring the highest abundance of SOX2/TBXT cells which were located towards the bottom of the dish while elongated structures comprised a cavity and were exclusively composed of SOX2 positive cell.

Together, these results demonstrate that suppression of NODAL signalling is necessary for the specification of NMPs and that adequate NODAL suppression and colony size combinatorially dictate uniaxial growth on micropatterns.

### 4 Small variations in CHIR dosage leads to radically distinct cell fate patterning outcomes

None of the tested conditions so far allowed us to find NOTO+ cells using FISH staining at 48h (not shown) raising the question of what other cues would be needed for NotoPs emergence. To gain further insights into the mechanisms of cell fate patterning on micropatterns upon CHIR exposure, and perhaps identify clues about the missing conditions for NotoPs emergence, we decided to test a range of CHIR concentrations and monitor markers of cell fates 48h post-induction via quantitative immunofluorescence (FIG 4A,B) and Nanostring analysis (Fig 4C). In the absence of CHIR, all the cells remained OCT4/SOX2 double positive and negative for TBXT, SOX17 and FOXA2 (FIG 4A, B and Sup Fig 4 A, B) showing that exogenous stimulation of the WNT pathway is required to initiate differentiation. More importantly, we found a clear dose-dependent effect of CHIR on the levels and spatial distributions of cell fate markers: The percentage of SOX2+ cells was negatively correlated to CHIR concentration while the percentage of TBXT+ cells increased proportionally, in line with the fact that TBXT is a known direct target of the WNT/β-catenin pathway (Arnold et al., 2000). On the other hand, SOX17 and FOXA2 did not correlate linearly with CHIR levels as these markers emerged at intermediate CHIR concentrations and were rare or absent at higher concentrations (Fig 4A and Sup Fig4 B). Instead, high doses of CHIR induced the expression of the lateral plate marker HAND1 (Sup Fig 4C). Our Nanostring analysis supported these results with a peak of endodermal markers at 1 and 2µM CHIR while colonies treated with the highest doses of CHIR expressed the LPM markers HAND1, TBX3 and the posterior marker CDX2 as well as HOX genes. These data suggest that endodermal cells are replaced by more posterior mesodermal fates at high CHIR concentration.

**Fig 4.**
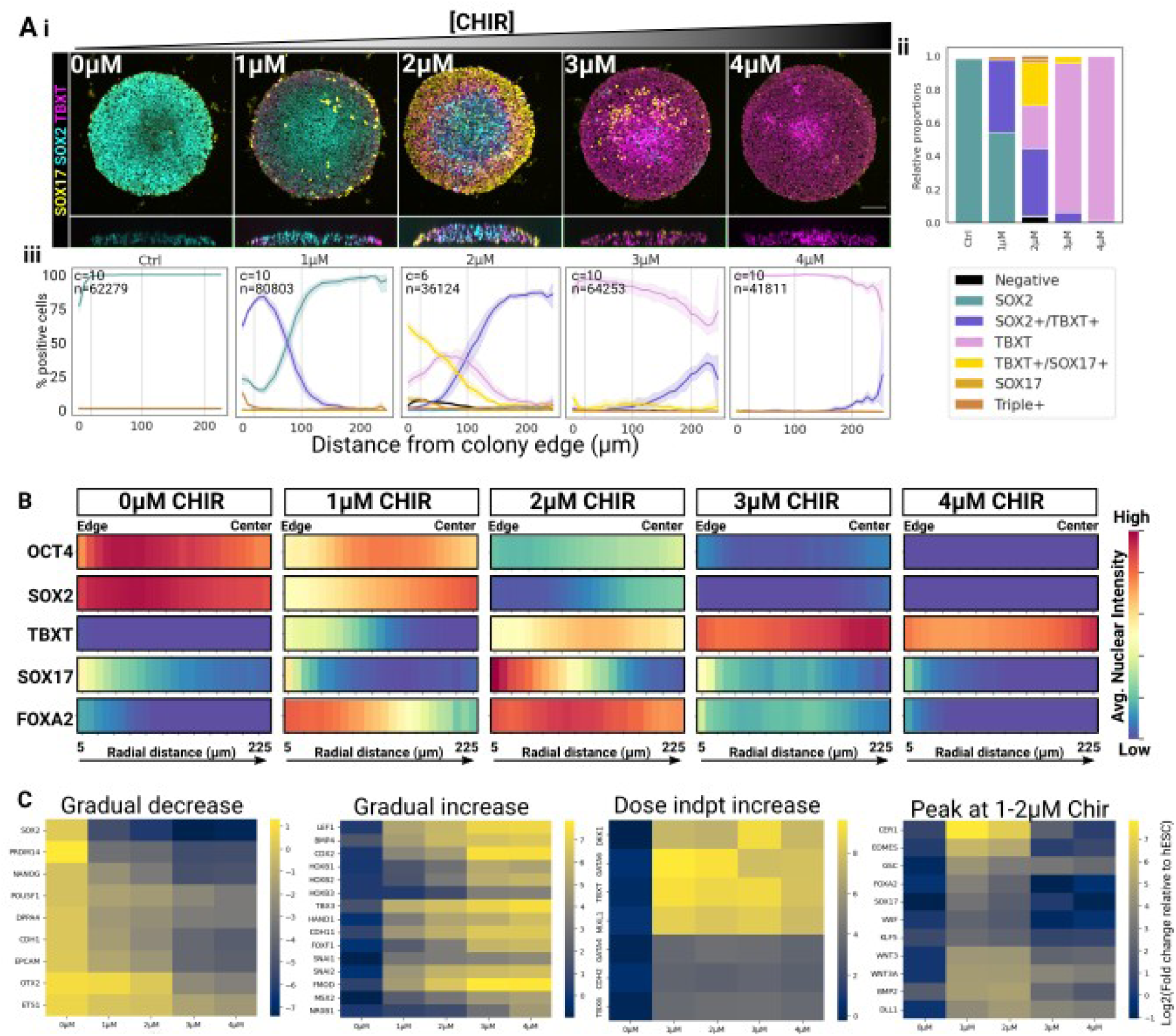
Cell fate patterning does not correlate linearly with CHIR dosage. **A** CHIR dose response in 500µm colonies fixed at 48h. (i) Images show the max projections of confocal z-stacks and a z-projection underneath. (ii) Stacked bar plot showing the relative proportions of individual cell populations for each CHIR concentration. (iii) Radial profiles of the percentage of each population. c: number of analysed colonies, n: number of analysed nuclei, shaded area: 95% confidence interval. **B** Heatmaps showing the radial profile of average nuclear marker intensity scaled between 0 and 1 on a per-marker basis. **C** Nanostring analysis of colonies grown in FGF and increasing doses of CHIR harvested at 48h. The colour scale corresponds to the log2 ratio of mRNA count in the sample relative to hESC.

Interestingly, when considering the spatial distribution of these markers, we observed a clear inward shift of the different cell populations when CHIR was increased (FIG 3A iii): TBXT was expressed at the periphery with 1µM CHIR and progressively expressed throughout the colony with increasing CHIR concentrations. In parallel, SOX2 was downregulated at the periphery with 2µM CHIR and higher concentrations were required to repress SOX2 in the centre. As a result, TBXT/SOX2 double positive cells were found at the periphery at 1µM CHIR but were progressively shifted and restricted to the centre, forming a smaller fraction of the total population with increasing CHIR concentration where TBXT single positive cells became predominant. Finally, SOX17 appeared restricted at the edge at 1µM CHIR, but spanned a 150 µm domain at 2µM, while at 3µM, the few SOX17+ cells present in the colony were located at around 100µm from the border (FIG 4A iii – yellow line). These observations further support the idea that CHIR-induced patterning is initiated from the periphery of the colony.

Together, our data show a complex, non-linear dose-dependent action of CHIR on cell fates, with intermediate doses of CHIR inducing endodermal cell fates predominantly, and higher doses inducing more posterior (FOXA2 negative) mesodermal fates.

### 5 CHIR dosage correlates with distinct downstream NODAL signalling dynamics

Our previous results raise the possibility of a non-linear effect of CHIR concentration on downstream signalling activity which would in turn define cell fate outcomes on micropatterns. We first checked the spatio-temporal dynamics of the canonical WNT pathway using the target gene LEF1 as a proxy (Fig 5A and Bi). We found that LEF1 expression increased monotonically over time at a rate proportional to CHIR concentration and we did not see any clear evidence of LEF1 spatial patterning along the colony radius. These data suggested that all the cells were all equally competent to respond to CHIR regardless of their position in the colony and that they did so proportionally to the dose of CHIR they received, suggesting that additional signals drive radial patterns of cell fates.

**Fig 5.**
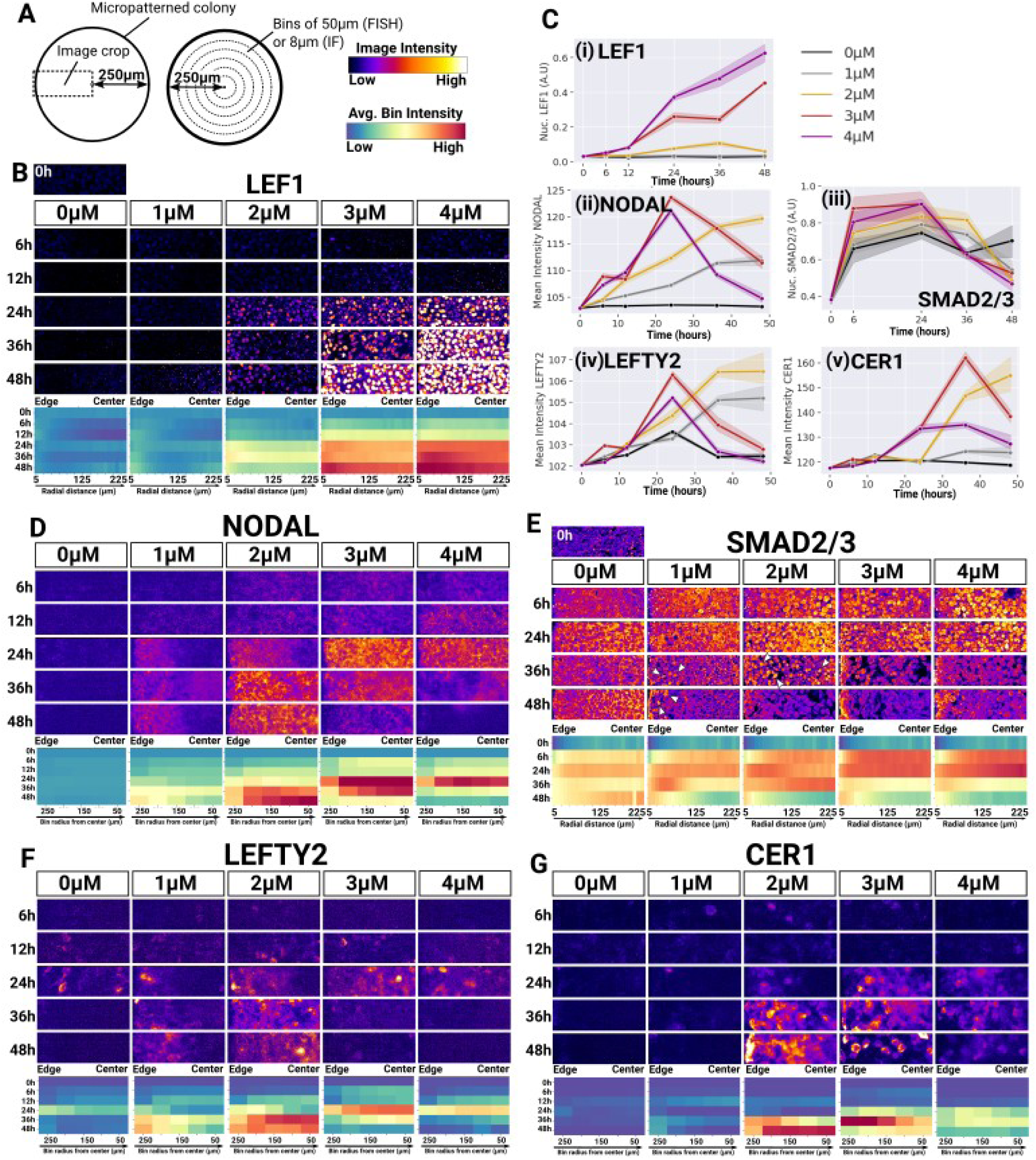
Small variations in CHIR dosage correlate with distinct downstream dynamics of NODAL signalling. **A** Cartoons describing the position of image crops shown in B and D-G relative to their respective colony (Left) and the organisation of the bins shown in heatmaps (Middle). Colour sales used in the figure are provided on the right. **B-G** Time course analysis of WNT and NODAL signalling in 500µm hESC colonies across CHIR concentrations. **B and E** show immunofluorescence analysis of nuclear LEF1 and nuclear SMAD2/3 levels respectively. Images show selected z-slice across a confocal z-stack. **D, F and G** show FISH analysis of NODAL, LEFTY2 and CER1 transcript levels. Images are 2D widefield images. For B, and D-G, the radial profiles of signal intensities over time are provided below images. **C** shows the temporal profiles or gene expression averaged across entire colonies. Lines indicate the mean average expression across colonies the shaded area indicate the 95% confidence interval. The figure is representative of two independent experiments.

Since we and others have found that CHIR induces NODAL signalling on micropatterns (Fig 2 - (Martyn et al., 2018; Massey et al., 2019), we next investigated the spatio-temporal dynamics of NODAL signalling across CHIR concentrations (Fig 5B-D). During the first 24h, the rate of increase in NODAL transcripts correlated with the dose of CHIR. However, while NODAL kept increasing after 24h with 1 and 2µM CHIR, NODAL expression rapidly decreased with 3µM CHIR and dropped even faster with 4µM (Fig 5A and B(ii)). Nuclear SMAD2/3 levels (the downstream signalling effector of the pathway) were consistent with the temporal profiles of NODAL expression. Indeed, nuclear SMAD2/3 first increased proportionally to CHIR concentrations at early time points and then dropped earlier at 3 and 4µM CHIR than at 1 and 2µM CHIR (Fig 5D and B(iii)) confirming the non-linear dependence of NODAL signalling to CHIR dosage.

Interestingly, when looking closely at the spatial pattern of NODAL signalling in the colonies, we observed that NODAL expression was evenly distributed during the first 12h at all CHIR concentrations (Fig 5C), consistently with the notion that all the cells responded to CHIR by activating NODAL expression. However, a radial gradient of NODAL expression starting from the periphery became apparent at 24h in colonies treated with 1 and 2µM CHIR. The NODAL expression domain expanded slightly at 1µM CHIR correlating with the positioning of a few cells with nuclear SMAD2/3 at 36h and 48h (Fig 5D). With 2µM CHIR, NODAL expression expanded to the centre as soon as 36h and became even stronger in the centre at 48h consistently with the presence of cells with nuclear SMAD2/3 across the entire colony at 36h. These results show that a radial gradient of NODAL expression (Fig 5C) precedes the emergence of endoderm at the periphery of colonies treated with 1µM and 2µM CHIR (Fig 2B and 4A). Such a wave of NODAL expression was not clearly apparent at 3µM and 4µM CHIR, perhaps because radial expression occurred too rapidly between 12h and 24h for our experiment to capture this process.

Interestingly, nuclear SMAD2/3 did not strictly follow NODAL expression. Furthermore, NODAL expression dynamics (FIG5 Bi) indicated the existence of a negative feedback that occurs faster with higher doses of CHIR. Inhibitors of the pathway include LEFTY2 and CER1 (Aykul et al., 2015). LEFTY2 expression followed closely the spatio-temporal dynamic of NODAL expression (FIG5 E and B (iv)) while CER1 harboured a similar pattern of expression albeit in a temporally delayed manner (FIG5 E and B (v)) supporting the idea that LEFTY2 and CER1 may indeed contribute to the temporal profile of NODAL expression and nuclear SMAD2/3. However, the fact that there exists a decoupling between the level of these inhibitors and the downregulation kinetic of NODAL (with 2µM CHIR for example) implies that other mechanisms are also at play and it will be interesting to elucidate this in the future.

All Together, our results demonstrate that even very small variations in CHIR concentration induce distinct NODAL signalling dynamics which correlate with cell fate outcomes. Importantly, a slow and sustained increase in NODAL expression together with a low expression of LEF1 was found in colonies which formed endoderm. On the other hand, a sharp transient expression of NODAL together with a strong increase in LEF1 was found in colonies where mesoderm was the most abundant. These experiments allowed us to gain insights into the mechanisms driving fate patterning in micropatterned colonies and begin to characterise a system that will be useful for more detailed mechanistic studies in the future.

### 6 Abrupt Nodal and BMP inhibition is required for the spontaneous emergence of NotoPs

We next turned to the question of how to further modify signalling in order to achieve NotoPs differentiation. Specification of NotoPs requires the cooperation of WNT and NODAL signals in the mouse embryo (Lickert et al., 2002; Vincent et al., 2003; Yamamoto et al., 2001). However, our previous results show that varying WNT activity alone is not sufficient to elicit NotoPs despite the consequences of WNT activity on downstream NODAL signalling. Therefore, we then asked whether endogenous BMP signalling may be preventing NotoPs emergence in our colonies. BMP signalling has been shown to inhibit specification of the notochord (Yasuo and Lemaire, 2001) and our Nanostring data showed that BMP2 and BMP4 are indeed expressed in our system (Fig 2 and Fig 4C), most likely downstream of NODAL signalling (Chhabra et al., 2019; Repina et al., 2023). We confirmed these results via FISH against BMP2 across CHIR concentrations at 48h (Sup Fig 5). Given these considerations, we hypothesised that the NODAL and BMP signalling dynamics established spontaneously within our colonies is inadequate for NotoPs emergence and that a tight exogenous control of TGFβ signalling together with sustained WNT activity is instead necessary.

To test this ideas, we inhibited NODAL and BMP signalling using small molecule inhibitors added either throughout differentiation or at 24h when a peak of NODAL expression was observed and when putative precursors of NotoPs may be present (TBXT/FOXA2 double positive cells were already present and SOX17+ cells were still absent at 24h). To monitor NotoPs emergence, we used FISH against the NotoP marker NOTO and against CHRD, a BMP inhibitor expressed in the APS and rapidly restricted to the node in mice (Bachiller et al., 2000).

As expected, we observed high NODAL expression and an absence of NOTO signal in the control (Fig 6A and B). Interestingly, we also found a broad domain of CHRD transcripts across the colony probably reflecting cells transiting through an early APS mesendoderm state. Treatment with 0.1µM of the BMP inhibitor LDN added at 24h had no effect on NOTO but led to an increase in CHRD and NODAL expression indicating that BMP signalling may be one of the factors negatively feeding back onto NODAL expression. More importantly, NODAL inhibition from 0h abolished NODAL and resulted in the presence of rare NOTO/CHRD double positive cells at the periphery of the colony, indicating that some NotoPs can be specified in this condition. Even more importantly, NODAL inhibition from 24h onwards induced a large domain of strong NOTO and CHRD co-expression localised within a 150µm distance from the colony edge. Addition of LDN at 24h further potentiated this effect confirming that BMP signalling inhibits NotoPs specification. NotoPs emergence was reproducible across all colonies within the experiment (Fig 7B) as well as with other cell lines (Sup Fig 6) confirming the robustness of these results. Furthermore, manipulating colony shape to maximise the proportion of cells experiencing edge effects such as lines of 200µm width enabled us to generate colonies with covered almost entirely of NOTO+ cells (Fig 7C)

**Fig 6.**
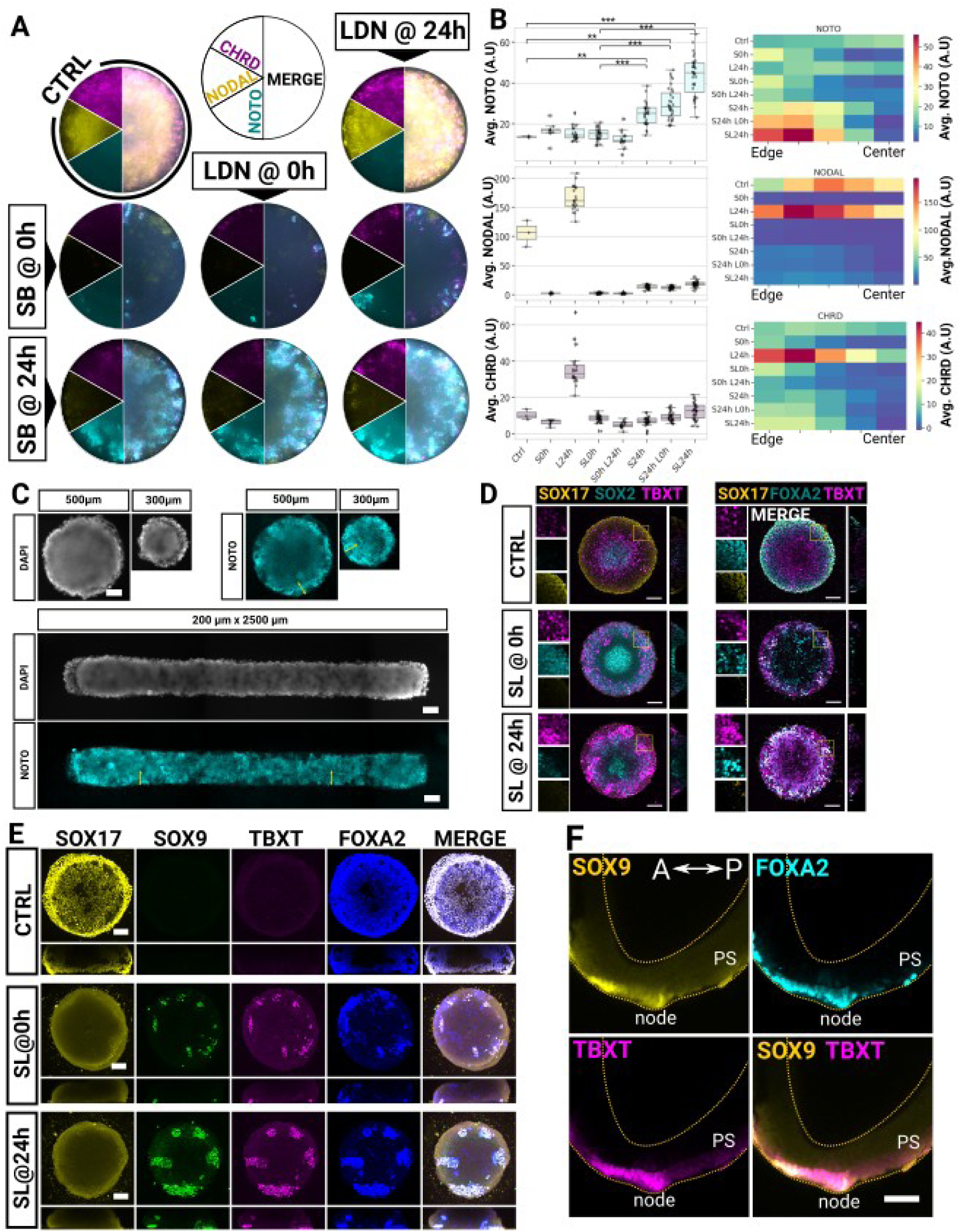
Timely TGFbeta inhibition is required for the spontaneous emergence of NotoPs on micropatterns. **A** FISH staining for NODAL, CHRD and NOTO in 500µm micropattern colonies induced with 2µM CHIR and 20ng/ml FGF2 and fixed at 48h post induction. Timed treatment with the Nodal or BMP inhibitors are indicated. **B** Shows box plots of average signal intensities in each individual colony (left) and heatmaps of binned signal intensities along the radial distance from the edge. Asterisks in box plots indicate the p-values of unpaired t-tests (** p<0.05; *** p<0.001). C Representative widefield images of micropatterned colonies stained via FISH at 48h with SL added at 24h. Yellow arrows indicate a 100µm domain from the edge. D Representative confocal z-stack max projections of colonies stained 2 days post-induction with CHIR and FGF and with or without SB and LDN (SL) treatment added at 0h or 24h. **E** Representative confocal z-stack max projections of colonies grown for 3 days in 2µM CHIR and 20ng/ml FGF containing medium and an additional 3 days in unsupplemented N2B27 with or without SL treatment added at 0h or 24h. **F** Confocal image showing a sagital view of a whole-mount immuno-stained early bud mouse embryo. Notice the SOX9 expression in the crown cells of the node and the nascent notochord. Scale bar: 100µm.

**Figure 7.**
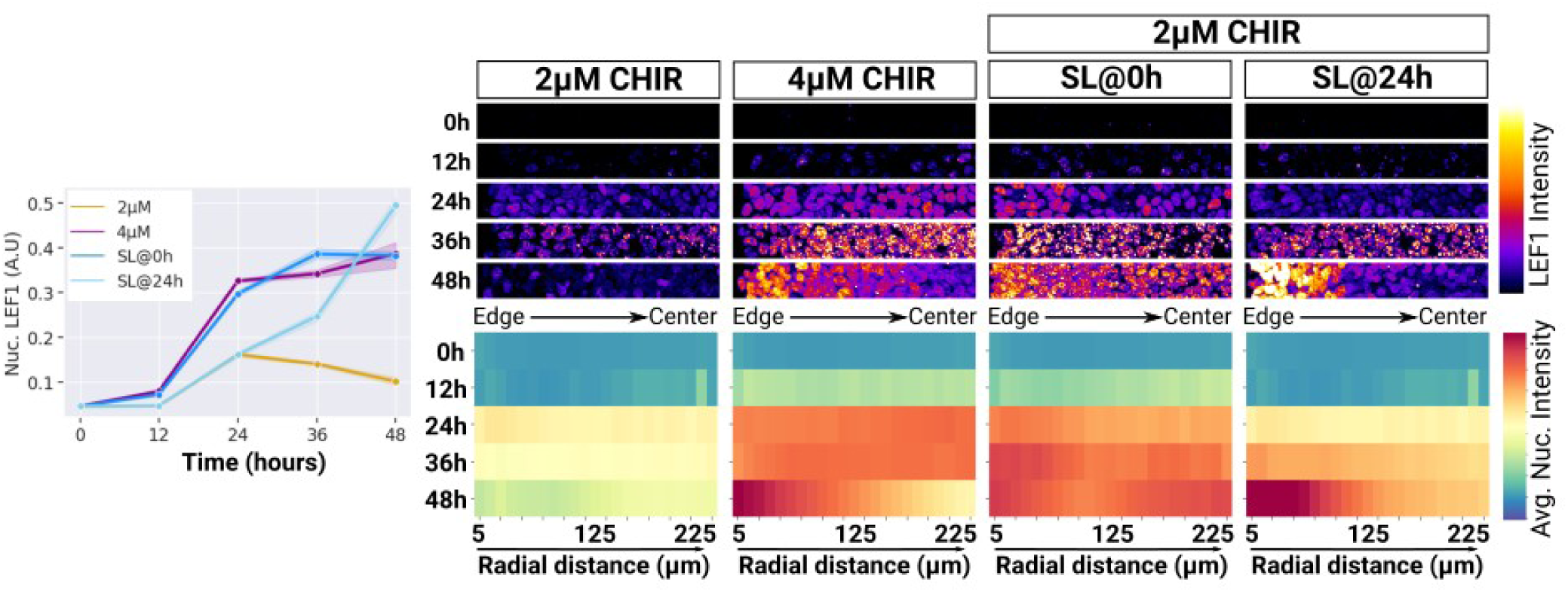
TGFbeta inhibition potentiates WNT signalling responsiveness in a domain that overlaps with NotoPs emergence. Time course analysis of LEF1 expression in 500µm hESC colonies. The line plot on the left shows the temporal profiles or average nuclear LEF1 intensity within 50µM of the colonies edges. The shaded area indicate the 95% confidence interval. Images on the right show representative image crops as indicated in Fig 5A. The radial profiles of signal intensities over time are provided as heatmaps below images.

Staining for SOX17, SOX2, TBXT and FOXA2 in these conditions at 48h (Fig 7D) showed that endoderm (FOXA2/SOX17 double positive cells) was eliminated from colonies treated with SB/LDN added at 24h and that instead the periphery of the colonies contained a large amount of TBXT+/ FOXA2+ and SOX17-cells. This result support the notion that FOXA2+/TBXT+ cells formed at 24h may be competent to form either endoderm or NotoPs depending on whether these cells experience sustained NODAL signalling or an abrupt inhibition respectively.

Finally, to determine whether the NOTO positive cells found at 48h were able to form notochord, we cultured the cells for three days with CHIR and FGF and then three more days in N2B27 alone with or without SB/LDN added at 0h or 24h (Fig 7E). We stained the cells for FOXA2, TBXT and SOX9 which are co-expressed in the node and the nascent notochord in early bud mouse embryos (Fig 7F) as well as SOX17 to distinguish cells that may have formed endoderm. While SOX9 and TBXT were absent from FOXA2/SOX17 co-expressing cells in the control colonies, we found a significant number of FOXA2, TBXT and SOX9 co-expressing cells when TGFbeta inhibitors were added in the medium. This number was further increased when SB and LDN were added at 24h.

All together our data provide evidence that delayed TGFbeta inhibition in micropattern colonies efficiently produce notochord-competent NotoPs.

### 7 TGFbeta inhibition potentiates WNT signalling response

Finally, we set out to understand the changes in signalling downstream of TGFbeta inhibition which might explain the specification of NotoPs. We used Nanostring to find differentially expressed genes in 48h colonies with or without the NODAL inhibitor SB added at 24h (Sup Fig 7).

Interestingly, we observed the upregulation of several NOTCH signalling related genes when NODAL was inhibited at 24h as well as genes indicating a more anterior identity such as GBX2 and HOXB2. Importantly, we also found a strong increase in the WNT target gene LEF1. To further elucidate how WNT signalling is impacted by TGFbeta inhibition and how this may be associated with NotoPs emergence, we performed a time course analysis of LEF1 expression in colonies treated with FGF and 2µM CHIR with or without SB and LDN added at 0h or 24h (Fig 7). We also included a 4µM CHIR condition to get a comparative measure of LEF1 levels in these colonies. Strikingly, when SB and LDN were added at 0h to colonies treated with 2µM CHIR, LEF1 intensity increased at the same rate as what was observed for 4µM CHIR treated colonies. When SB and LDN were added at 24h to 2µM CHIR treated colonies, LEF1 expression sharply increased to reach the levels observed in colonies treated with 4µM CHIR. These results are in line with our Nanostring data and demonstrate that TGFbeta inhibition strongly potentiates the cells responsiveness to canonical WNT signalling. Importantly, LEF1 intensity was the highest at the periphery of the colonies treated with SB and LDN added at 24h. This domain overlapped with the domain of NOTO+ cells suggesting that high WNT activity drives NotoPs emergence. Given that 4µM CHIR treatment generates a high level of WNT activity and a transient peak of NODAL expression (Fig 5) we wondered whether NotoPs could be obtained by simply inhibiting BMP signalling in 4µM colonies (Sup Fig 8). This, however did not result in any NOTO positive staining and neither did the addition of SB and LDN at 24h in these colonies. These results indicate that a tight control of the temporal profiles of WNT and TGFbeta signalling is needed for NotoPs emergence.

Together our data clarify how WNT and TGFbeta signalling cooperate in order to specify the notochordal lineage. While WNT and TGFbeta signalling are initially necessary to induce an early APS cell state, a timely and abrupt inhibition of TGFbeta signalling is necessary to maximise WNT signalling response and define NotoPs.

## Discussion

The series of lineage restrictions that occur in the APS during gastrulation remain challenging to investigate *in vivo* and this is especially true in a human context. Here, using spatial confinement with micropatterns, we were able to direct the development of hESC into reproducible radial patterns of all the APS cell fates including NotoPs (Colombier et al., 2020). This system enabled us to gain insights into 1) the mechanisms that regulate temporal patterns of signalling cues and 2) to delineate the signalling sequences distinguishing APS cell fates from one another (Figure 8).

**Figure 8.**
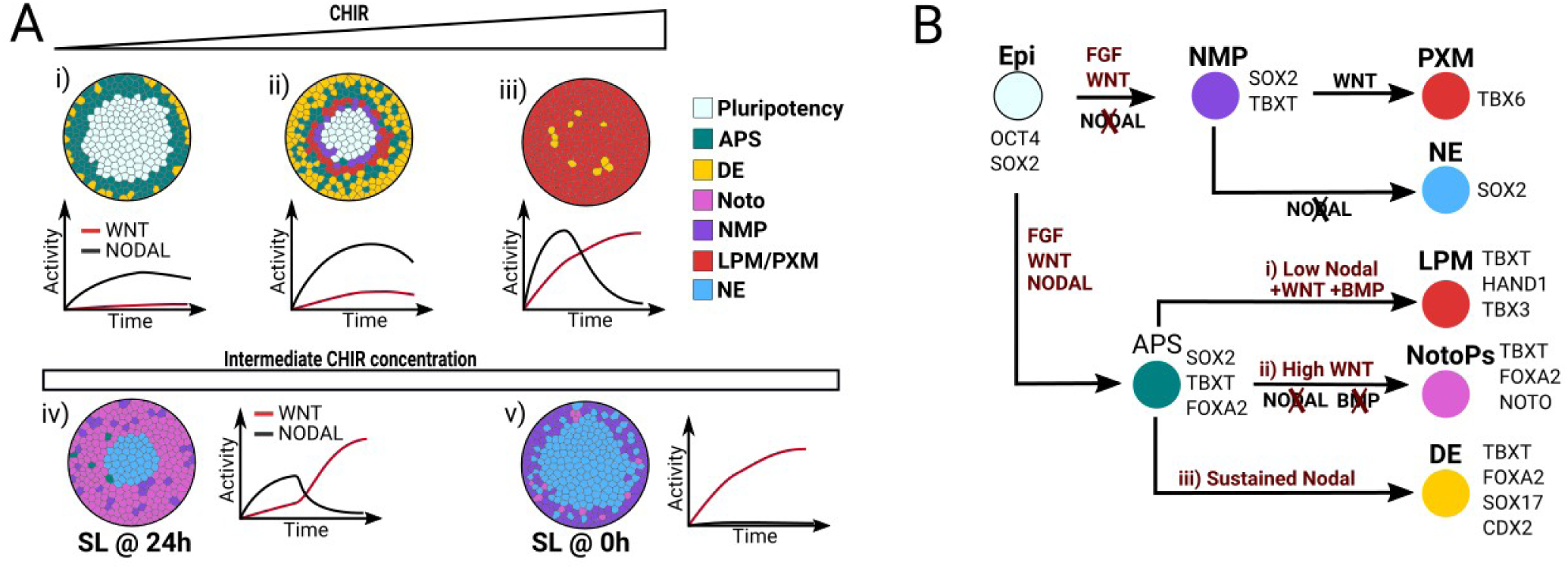
Graphical summary of cell fate outcomes and associated signalling dynamics. **A** Diagram summarising the spatial organisation of cell fates at 48h on micropatterns when treated with 20ng/ml FGF and with 1µM CHIR (i), 2µM CHIR (ii, iv, v), 3µM CHIR (iii) and SB and LDN added at 24h (iv) or 0h (v). THE WNT and NODAL signalling profiles associated with these patterns of differentiation is indicated as a line plot underneath each colony. **B** shows the putative hierarchy of cell fate lineages and the signalling conditions defining lineage restrictions.

Our first experiment showed that cells on micropatterns treated with 2µM CHIR and 20ng/ml FGF2 predominantly underwent endodermal differentiation on micropatterns (Fig 1B). This was at first surprising to us given that the same concentrations of CHIR and FGF in 2D monolayer cultures normally lead to homogenous NMP differentiation (Fig 1B). This also raised the question of what makes the cells on micropatterns adopt a distinct fate than in monolayer cultures.

Previous work using micropatterns and BMP or WNT ligands as the triggers of differentiation have shown that cells located at the colony boundary respond more efficiently to signalling molecules than cells in the centre due to a loss of epithelial integrity on the colony edges (Etoc et al., 2016; Legier et al., 2023; Martyn et al., 2018; Martyn et al., 2019; Warmflash et al., 2014). Our data indicate that a similar boundary-driven mechanism is likely taking place here as well given that the size of the peripheral differentiation domain remained constant with increasing colony sizes (Sup Fig 2). In our case however, the cells were treated with CHIR, a cell-permeable molecule that bypasses ligand/receptor interactions and is therefore unlikely to be affected by a loss of epithelial integrity. Our LEF1 staining showed that all the cells were indeed equally responsive to CHIR (Figure 5A). Nevertheless differential responsiveness to ligands may matter when secondary signals downstream of CHIR are secreted. Our data support this idea as NODAL expression was initiated throughout the colony at early time points (as a result of equal responsiveness to CHIR) before a wave of NODAL expression travelling from the periphery towards the centre became apparent (Figure 5D). Such a wave, which was also described in other systems (Heemskerk et al., 2019; Liu et al., 2022; Martyn et al., 2019) may be the result of the positive self-regulation of NODAL (Liu et al., 2022) together with a high responsiveness to NODAL at the periphery. As NODAL signalling is a notorious driver of the endodermal lineage (Nowotschin et al., 2019) and NODAL inhibition prevents endoderm emergence (Fig 3), we conclude that confinement potentiates NODAL signalling downstream of CHIR which in turn drives endodermal differentiation on micropatterns.

Our data are in agreement with and complementary to a recent preprint showing that treatment of micropatterned colonies with WNT3a and FGF8 ligands generates a similar spatio-temporal pattern of NODAL expression (Fig5J of (Ortiz-Salazar et al., 2024)). Both systems ultimately lead to the formation of reproducible and self-organised 3-dimensional structures composed of definitive endoderm with a core of epiblast cells as shown in Fig 1F. These micropattern systems therefore provide a powerful platform - complementary to other 3D human in vitro models (Moris et al., 2020; Vianello and Lutolf, 2020) - to further understand endoderm specification and morphogenesis in a human context.

Importantly, we found that endoderm only emerges on micropatterns in a very small CHIR concentration range around 2µM. Our CHIR dose-response experiment (Fig 4) was particularly revealing and helped us to delineate the signalling sequences that segregate individual APS cell fates from one another (summarised in Fig 8). We show that even very small variations in CHIR concentrations radically change the proportion of endodermal and mesodermal cell fates (Fig 4 and Fig 8) and that this effect was mediated by a non linear relationship between CHIR concentration and the downstream dynamics of NODAL expression (Fig5C-E). 1 and 2µM CHIR induced a progressive and sustained increase in NODAL production favouring the endodermal lineage while higher CHIR concentrations resulted in a sharp increase followed by a rapid reduction of NODAL expression which correlated with an abundance of mesoderm (Fig 4, 5 and 8). It is useful to compare these findings with the results from Ortiz-salazar et al. These authors reported that a 10-fold increase in WNT3a ligand concentration did not alter endoderm emergence while CHIR (10µM) induced mesoderm differentiation. Overall, our data are in agreement and complementary and may identify the range of CHIR concentrations that relate to the physiological, ligand-based activation of the pathway.

Our findings also suggest the existence of a negative feedback on NODAL expression when NODAL reaches high levels of expression (Fig 5B). Secreted antagonists of the pathway, LEFTY2 and CER1 may contribute to the downregulation of NODAL at high CHIR concentrations but are unlikely to be the sole negative regulators of NODAL given the decoupling between the level of expression of these antagonists and the kinetic of NODAL downregulation (Fig 5). Furthermore, we found heterogeneous levels of nuclear SMAD2/3 in individual colonies with domains of NODAL expression that did not overlap necessarily with nuclear SMAD2/3 (Fig 5D and E). One possible reason include an antagonism between NODAL and BMP. Indeed, BMP has been shown to antagonise NODAL in other contexts (see for example (Pereira et al., 2012)) and we found here that higher CHIR dosages correlated with higher levels of BMP2 transcripts (Sup Fig 5). It will be interesting to use our micropattern system to elucidate this question in the future.

Most importantly, our data sheds some light on how distinct signalling regimes are established and associated with individual cell fates (Fig 8B):

1. We have found that NMP specification requires an environment where NODAL activity is maintained at a minimum (Fig 3). This idea is in agreement with a recent report showing that the maintenance of NMPs *in vitro* requires NODAL inhibition (Kelle et al., 2024). This notion is also compatible with work in mouse embryos showing that NMPs appear later than endoderm, at a time when NODAL signalling activity starts to decrease (Lawson et al., 1991). Furthermore, neighbouring NotoPs - as a source of NODAL inhibitors - may protect NMPs from advert differentiation during axial elongation. Indeed a release of NODAL inhibition might explain why NotoPs ablation results in early termination of axial elongation (Abdelkhalek et al., 2004; McLaren and Steventon, 2021; Saunders et al., 2024; Wymeersch et al., 2019).
2. Our observations also show that a proportion of the cells differentiate to lateral plate mesoderm (Fig 1F), most likely under the influence of endogenous BMP signalling (Heemskerk et al., 2019). This is perhaps not surprising as fate mapping experiments have shown that cardiac mesoderm arise directly adjacent and posterior to the definitive endoderm (Lawson et al., 1991; Tam et al., 1997) and lineage tracing using FOXA2-cre mouse lines have demonstrated that a proportion of FOXA2 expressing cells in the streak are fated to form cardiac ventricles (Bardot et al., 2017). It will be interesting in the future to confirm the lineage hierarchy and the identity of the HAND1+ cells emerging in our system.
3. Finally, our manipulations of NODAL and BMP signalling has enabled us to identify the signalling requirements for the specification of the notochord lineage (Fig 6A and 8B) and unveiled a signalling cross-talk where NODAL signalling dampens WNT activity (Fig 7). We propose a model where initial WNT and NODAL signalling combinatorially specify epiblast cells to an early multipotent APS state (co-expressing TBXT and FOXA2) which can commit to either endoderm or notochord. While prolonged NODAL exposure keeps WNT activity low and drives endoderm differentiation, a sharp inhibition of NODAL signalling potentiates WNT activity and redirects the cells to the NotoP fate (Fig 8B). Our results and model are consistent with observations in the Xenopus where a sudden drop of p-Smad2 correlates with the emergence of the notochord (Schohl and Fagotto, 2002) but it will be important to test our model in an *in vivo* setting and determine mechanisms that are evolutionary conserved or that differ in the human.

Reassuringly, our data are reproducible across multiple cell lines (Sup Fig 6) and consistent with a recent study reporting a similar requirement for delayed TGFbeta signalling inhibition for the derivation of NotoPs on micropatterns (Rito et al., 2023). These authors also report a cross-talk between hippo signalling and FGF activity at the colony boundary overlapping with the domain of NotoPs emergence.

Our data showing that TGFbeta inhibition potentiates WNT activity in this domain nicely complement these findings. It will be important in the future to elucidate whether and how the specific mechanical and biochemical environment defined at the colony boundaries *in vitro* are defined inside the embryo.

## Conclusion

NotoPs are regarded as a promising cell type for drug discovery or cell therapy and much remains to be learned about the healthy and pathological development of the notochord. Encouragingly, recent evidence suggest that NotoPs persist as a small transcriptionally stable population throughout axial elongation (Wymeersch et al., 2019) making it likely that NotoPs may be expandable in culture if the correct conditions can be identified. Our work together with the recent work from Rito and colleagues (Rito et al., 2023) provides insights that will inform the development of reliable NotoPs derivation protocols. It will be interesting to test if a fully functional node can be reproduced *in vitro* and determine what other signalling cues define the proportions and maturation of individual node sub-populations in the future.

## Materials and Methods

### Cell culture

Experiments were conducted with the MasterShef7 hESC line obtained from the University of Sheffield, the RC17 hESC line from the University of Edinburgh (De Sousa et al) and the NAS2 hIPSC line from the Kunath lab (Devine et al). Approval for the use of hESC was granted by the steering committee for the UK Stem Cell bank and for the Use of Stem Cell Lines. Cell lines were propagated at 37°C and 5% CO_2_ in mTSER Plus medium (100-0276, Stemcell Technologiess) on Geltrex (A1413302, Life Technologies) coated 6-well plates (3516, Corning Incorporated). Wells were coated for 30 minutes at 37°C using a 100µg/mL Geltrex solution diluted in Magnesium and Calcium containing DPBS (14080-048, GIBCO). Passaging was performed every 2 to 3 days using Accutase (00-4555-56, ThermoFisher Scientific). The cells were Mycoplasma tested prior to running experiments. Only cells with a passage number below 30 were used for all experiments.

### Micropattern fabrication

Micropatterning was done in Ibidi 8-well µSlides (IB-80826, Ibidi) with a PRIMO bioengineering plateform (Alveole) mounted on a Nikon Ti2 Widefield microscope using a protocol adapted from Alveole: The surface of ibidi slides was first passivated as follows: slides were plasma treated (Harrick Plasma Cleaner) for 1 min 30 seconds at high intensity in a 200 mTorr vacuum and then incubated at room temperature for 1h with 200µL/well of 200 µg/mL Poly-D-Lysine (A-003-E, Merck Millipore) diluted in 0.1M HEPES pH 8.4 (H3784-25G, Sigma-Aldrich). The wells were washed twice with ddH_2_O and once with 0.1M HEPES pH 8.4 and then incubated for 1h in the dark with 125µL/well of 80mg/mL mPEG-SVA (MPEG-SVA-5000, Laysan Bio) freshly dissolved in 0.1M HEPES pH 8.4. Slides were then washed profusely with ddH_2_O, air dried and stored at 4°C until further processing (1 week maximum). The PRIMO insolation step was next performed less than one day prior to plating the cells: passivated wells were covered with 8µL PLPP gel (Cairn Research, 1µL PLPP gel/well diluted with 7µL 70% Ethanol) in the dark and left to dry for ∼30min at room temperature. Slides were then insolated with PRIMO through a 20X lens with a dose of 50mJ/cm^2^. All micropattern shapes were designed in Inkscape and converted to binary tiff files in ImageJ prior to loading in the Alveole Leonardo software. After insolation, PLPP gel was removed with 3 ddH2O washes and the slides were air dried and stored at 4°C until use.

### Culture on micropatterns

Micropatterned Ibidi slides were first rehydrated for 5 min in Magnesium and Calcium containing DPBS (14080-048, GIBCO; thereafter DPBS++). Matrix coating was then performed by incubating the wells at room temperature for 30 min on a rocker with a mixture of 40µg/mL rhVitronectin-N (A14700, ThermoFisher) and 10µg/mL rhLaminin521 (A29249, GIBCO) diluted in DPBS++. Wells were washed 3 times with DPBS++ and left in the last wash whilst preparing the cells for seeding to ensure that the wells were not left to dry. For seeding, 80% confluent cells were dissociated to single cells with Accutase, and resuspended in seeding medium composed of mTESR Plus supplemented with 10µM Y-27632 (1254, Tocris Bio-Techne) and 1:100 Penicillin/Streptomycin (10,000U/mL pen, 10,000mg/mL strep; 15140-122, Invitrogen). Cells were plated onto micropatterns at a density of 150 000 cells/well in 250µL of seeding medium. The cells were left to adhere for 3h at 37°C. After attachment, the excess of cells was removed by gentle pipetting and applying fresh seeding medium. The cells were left to settle and cover the patterns overnight until induction of differentiation the next morning. Cells were washed once in N2B27 to remove traces of growth factors present in seeding medium. Differentiation was then induced using N2B27 medium supplemented with Penicillin/Streptomycin (1:100), CHIR 99021 at concentrations indicated in figures legends (4423/10, Tocris Bio-Techne), 20ng/ml human bFGF (PHG6015, ThermoFisher Scientific). SB 431542 (1614, Tocris Bio-Techne) was used at 10µM unless specified otherwise and LDN 193189 (72147, StemCell Technologies) at 0.1µM.

### Immunofluorescence

Mouse embryos were staged according to (Downs and Davies, 1993) and stained as described previously in (Wong, 2021). Cells grown on micropatterns were fixed with 4% PFA (CHE2036, Scientific Laboratory Supplies), washed 3 times with a solution of PBS and 0.1% Triton X-100 (A16046, Alfa Aesar; hereafter PBST) and left overnight at 4°C in blocking solution containing 3% Donkey Serum (D9663, Merck) and 0.03% Sodium Azide (40-2000-01, Severn Biotech Ltd) in PBST. All primary antibodies were incubated overnight at 4°C and secondary antibodies at room temperature for 3h. All washing steps were performed in PBST. Some co-staining required the use of primary antibodies raised in the same species. In these cases, the staining was performed with either pre-conjugated antibodies only, or sequentially using a non-conjugated antibody first, its corresponding secondary antibody next, followed by a blocking step using species-specific serum (3% Goat Serum (G9023-10ML, Sigma) or 3% Rabbit Serum (R9133, Merck)) and finally applying the conjugated antibody. Slide were washed 3 times in PBST and kept sealed at 4°C until imaging.

| Primary Antibody | Cat. no, supplier | Dilution |
| --- | --- | --- |
| SOX2 | 14-9811-82, ThermoFisher Scientific | 1:200 |
| TBX6 | AF4744, R&D Systems | 1:400 |
| PAX6 | 561462, BD Bioscience | 1:300 |
| TBXT | NL2085R Conjugated, R&D Systems | 1:300 |
| SOX17 | AF1924, R&D Systems | 1:200 |
| CDX2 | Ab76541, Abcam | 1:200 |
| OCT4 | AP3724A, Abcepta | 1:200 |
| FOXA2 | 8186, Cell Signalling | 1:400 |
| SMAD2/3 | 610842, BD Bioscience | 1:100 |
| SOX9 | Ab185230 Conjugated, Abcam | 1:250 |
| CDH1 | SIGMA, MABT26 | 1:100 |
| CDH2 | BD Bioscience, 610182 | 1:200 |

| Secondary Antibody | Cat. no, supplier | Dilution |
| --- | --- | --- |
| Donkey-Anti-Goat-405 | Abcam, ab175665 | 1:1000 |
| Donkey-Anti-Mouse-405 | Abcam, ab175658 | 1:1000 |
| Donkey-Anti-Rat-488 | Abcam, ab150153 | 1:1000 |
| Donkey-Anti-Mouse-488 | Abcam, ab150105 | 1:1000 |
| Donkey-Anti-Rabbit-A568 | Abcam, ab175470 | 1:1000 |
| Donkey-Anti-Mouse-568 | Abcam, ab175472 | 1:1000 |
| Donkey-Anti-Rabbit-647 | Abcam, ab150075 | 1:1000 |
| Donkey-Anti-Rat-647 | Abcam, ab150155 | 1:1000 |
| Donkey-Anti-Mouse-647 | Abcam, ab150107 | 1:1000 |

### Fluorescent In Situ Hybridization

Branched DNA FISH was performed using the viewRNA Cell Plus assay from ThermoFisher Scientific (88-19000-99) according to the manufacturer’s protocol. FISH probes used in this study are listed in the table below. Slides were imaged on the same day with a Nikon TiE widefield inverted microscope and a dry 20X lens.

| FISH Probe-Channel | Cat.no |
| --- | --- |
| NOTO-488 | VA4-3101015-VC |
| CHRD-546 | VA1-3005411-VC |
| NODAL-647 | VA6-3167894-VC |
| LEFTY2-546 | VA1-3004742-VC |
| DKK1-488 | VA4-3082344-VC |
| CER1-488 | VA4-3089614-VC |
| BMP2-647 | VA6-3168104-VC |

### Immunofluorescence Imaging and Image analysis

Embryos were imaged in PBST using a Leica TCS SP8 Confocal and a 25X water immersion lens. All images were annotated and contrast-adjusted using FIJI (Schindelin et al., 2012). Micropatterned colonies were imaged with an Opera Phenix Plus (Perkin Elmer). Ibidi slides contained around 64 colonies per well, out of which ∼20 colonies were selected for analysis in each experiment. To ensure an unbiased sampling of the colonies, the slides were first fully scanned with a 10x lens to generate overview images of the LMBR (nuclear envelope marker) signal only. These images were then processed with an automated pipeline in the harmony software (Perkin Elmer). This pipeline rejected colonies with an unexpected area or roundness and then randomly sampled 20 colonies from the pool of valid colonies. Sampled colonies were next imaged with a 20X lens to generate 3D multichannel z-stacks with voxel size of 0.59 x 0.59 x 1µm. Opera images were then exported as Tiff files for further analysis. Nuclear segmentation was performed on the LMBR signal as described previously (Blin et al., 2019). Raw images and nuclear masks were imported into Pickcells (https://pickcellslab.frama.io/docs/) to compute nuclear features including 3D spatial coordinates and average intensities in all fluorescence channels. The tsv file created in PickCells was then analysed in python. Our Jupyter Notebooks and data files are available in our Gitlab repository (https://framagit.org/pickcellslab/data/2023_haxioms).

### FISH Imaging and Image analysis

Micropatterened colonies were imaged on an inverted widefield microscope (Nikon Eclipse Ti) with a long distance 10x lens to image entire wells (8mm x 8mm). Images were were then segmented using CellPose 2.0 (Stringer et al., 2021) to identify individual colonies. We used the livecell model as a starting point and manually adjusted the model according to the CellPose tutorial using at least 5 different images to ensure accurate segmentation across our conditions. Images were scaled down for the segmentation process to save on computational resources and scaled back up to their original sizes using ImageJ prior to loading into PickCells for further analysis. PickCells was used to compute average intensities in each channel within each colony, as well as background intensity in the immediate vicinity of the colony (bounding box intensity – colony intensity) and radial intensities as described in Fig 5. Data were next analysed in python. The colonies were selected based on their compactness and surface area, to avoid including abnormally sized and shaped colonies (e.g. partially broken colonies). The background intensity was subtracted from the average and radial colony intensities and plotted in python.

### Nanostring analysis

RNA samples were prepared using an Absolutely RNA microprep kit (cat.no 400805, Agilent technologies) and Nanostring profiling was performed using nCounter technology as per the manufacturer’s instructions. We used a panel of probes consisting of the 780 genes included in the standard human embryonic stem cell gene panel together with 30 additional custom probes (all genes are listed in Sup Table 1). Normalisation of raw data was accomplished in the Nanostring dedicated nCounter software. Next, raw counts were imported in R (R Core Team, 2013) and analysed with the Bioconductor package moanin (Varoquaux and Purdom, 2020). We first applied an initial cut-off to filter out all the genes where the max count was below 100. The data was then log2 transformed. We kept only the top 50% most variable genes based on the median absolute deviation (mad) metric over time. A spline was then fitted onto each individual gene profile and we grouped genes into 7 clusters using kmeans clustering on the parameters of the fitted splines to obtain the heatmaps shown in Fig 2A. R scripts and data are available in our Gitlab repository (https://framagit.org/pickcellslab/data/2023_haxioms).

### Mouse husbandry

Mouse work was carried out under the UK Home Office project license PPL PEEC9E359, approved by the University of Edinburgh Animal Welfare and Ethical Review Panel and in compliance with the Animals (Scientific Procedures) Act 1986. Mice used were of C57Bl6/J strain background. The mice were kept under a 12 h light/dark cycle, and the embryo age was denoted day 0 on the midpoint of the dark cycle the day the plug was found.

## Funding

This work was funded by a BBSRC project grant to GB ref:BB/W002310/1, a BBSRC Alert equipment grant to GB ref: BB/T017961/1 and a WT ISSF3 award to GB ref: IS3-R1.16 19/20. MRG is funded by a BBSRC EASTBIO studentship. EO was funded by a BSDB Gurdon studentship.

## Acknowledgements

We thank Tilo Kunath for the provision of RC17 and NAS2 cell lines. We also thank Sally Lowell, Anestis Tsakiridis and Anne Camus for their critical review of the manuscript. We also thank Val Wilson for helping us with mouse embryos. We thank Justyna Cholewa Waclaw and Matthieu Vermeren for their help with imaging and Alison Munro for running the Nanostring samples.

## Notes

### Competing Interest Statement

The authors have declared no competing interest.

### Summary of Updates

* Clearer images and annotations in Figure 1 to show the evolution of the cells on micropatterns * New data showing axial elongation and NMP emergence in Figure 3 * Added nanostring data in Fig 4 * More data to document signalling dynamics on micropatterns in Figure 6 * Added FISH for NOTO on micropatterns of different shapes and sizes in Fig 6 and better annotations of the embryo image. * New data to better document WNT and NODAL signalling cross-talk and dynamic in Fig 7 * Better summary diagram in Fig 8. * Added supplementary figures to show that the data are reproducible with other cell lines

