## Supplementary figures for "In vitro modelling of anterior primitive streak patterning with human pluripotent stem cells identifies the path to notochord progenitors"

**Supplementary Figure 1 Cell fate patterning is reproducible across 3 different cell lines**

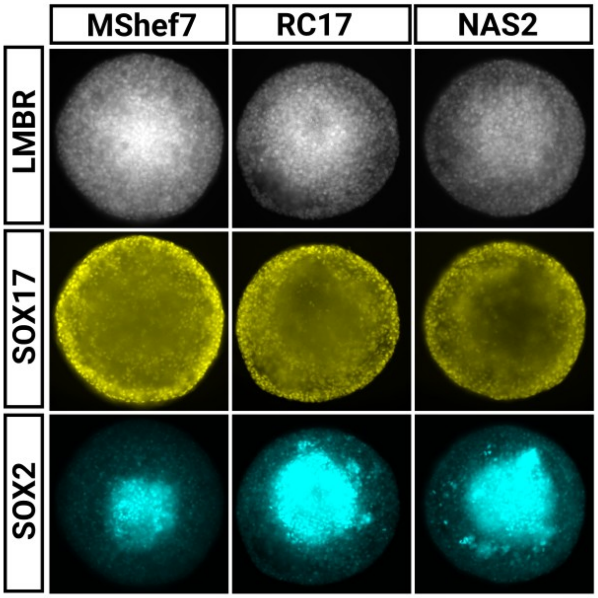


### Supplementary figure 1

Representative widefield images of hPSC colonies treated with 2µM CHIR and 20ng/ml FGF and stained at 48h post induction with the enodermal marker SOX17 and the pluripotency marker SOX2. The patterning of endodermal cells is reproducible across the 2 hESC tested (MShef7 and RC17) and one iPSC cell line (NAS2)

**Sup Fig 2 Cell fate patterning does not scale with colony size**

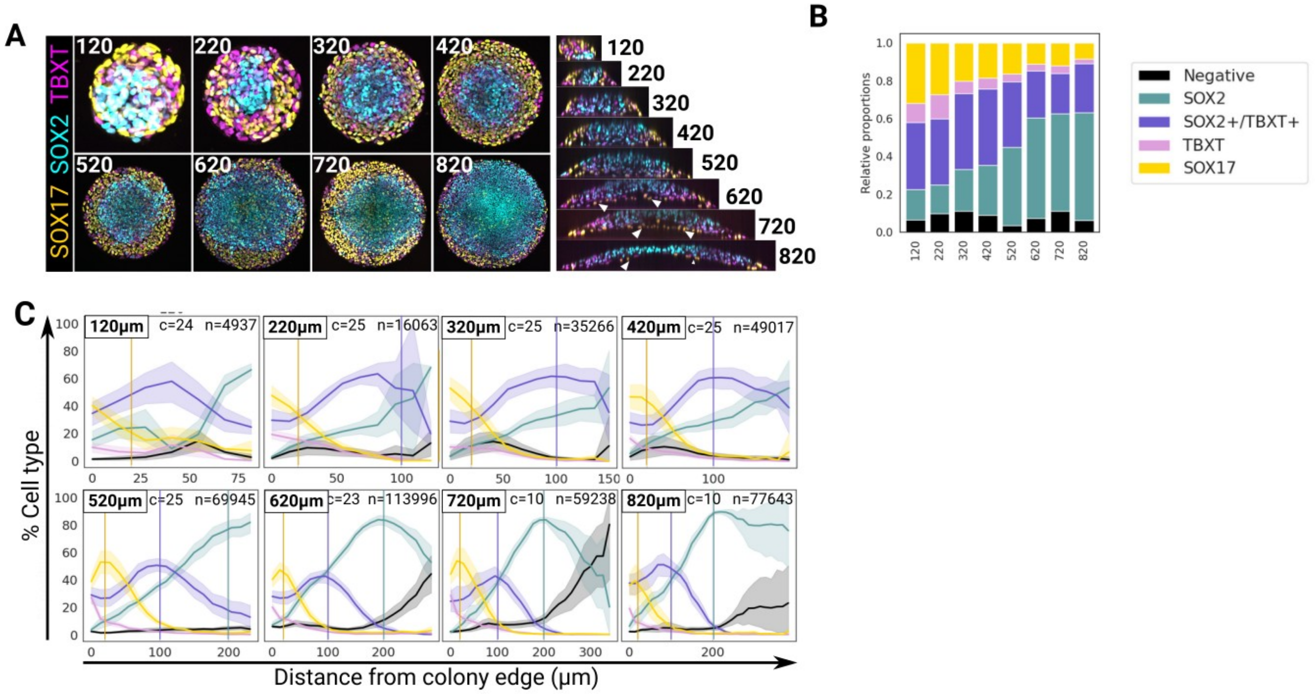


**Supplementary figure 2**

**A** Confocal max projections of micropatterned colonies of increasing diameters (indicated in the top left corner in µm) and stained for SOX17, SOX2 and TBXT. The corresponding Z projections are shown on the right with white arrowheads pointing at SOX17+ cells lining the central domain at the bottom of the colony. **B** Stacked bar plot showing the relative proportions of individual cell populations found for each colony diameter. The number of colonies and nuclei analysed for this plot are the same as in iii. **C** Line plots showing the mean proportion of each cell population as a function of the radial distance from the edge of the colonies. c: number of analysed colonies, n: number of analysed nuclei, shaded area: 95% confidence interval.

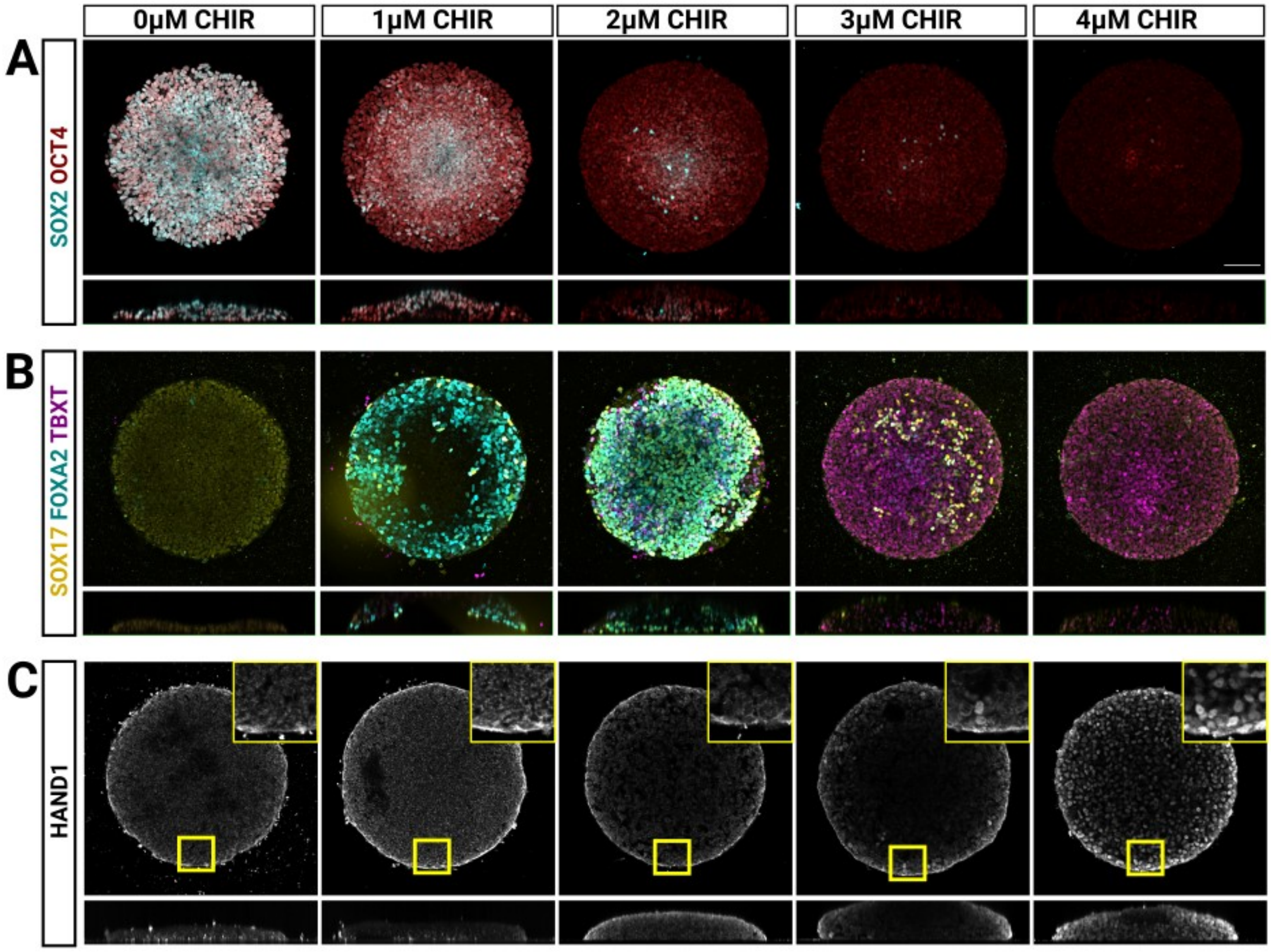


**Supplementary figure 3**

CHIR dose-response in 500μm colonies 48h post-induction. **A and B** include representative max projections of confocal z-stacks while **C** shows a selected z-slice across a representative confocal z-stack to better show the nuclear localisation of HAND1

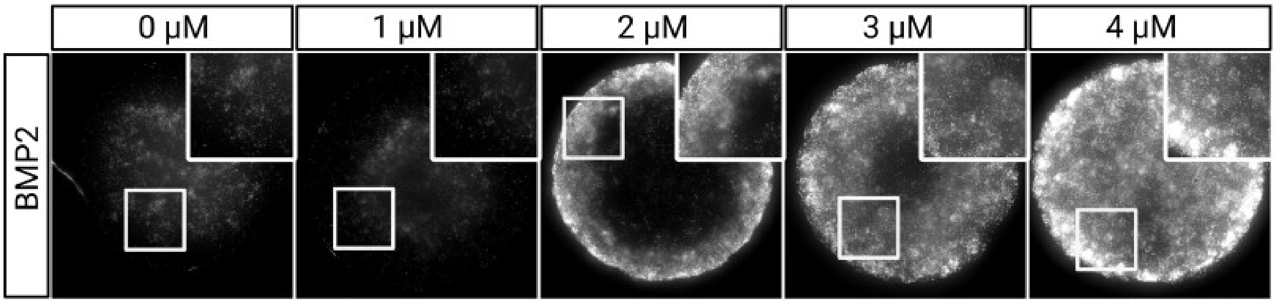


**Supplementary figure 4**

Representative widefield images of FISH staining against BMP2 transcripts in 500µm colonies 48h post-induction treated with 20ng/ml FGF2 and a range of CHIR concentrations indicated at the top of the images.

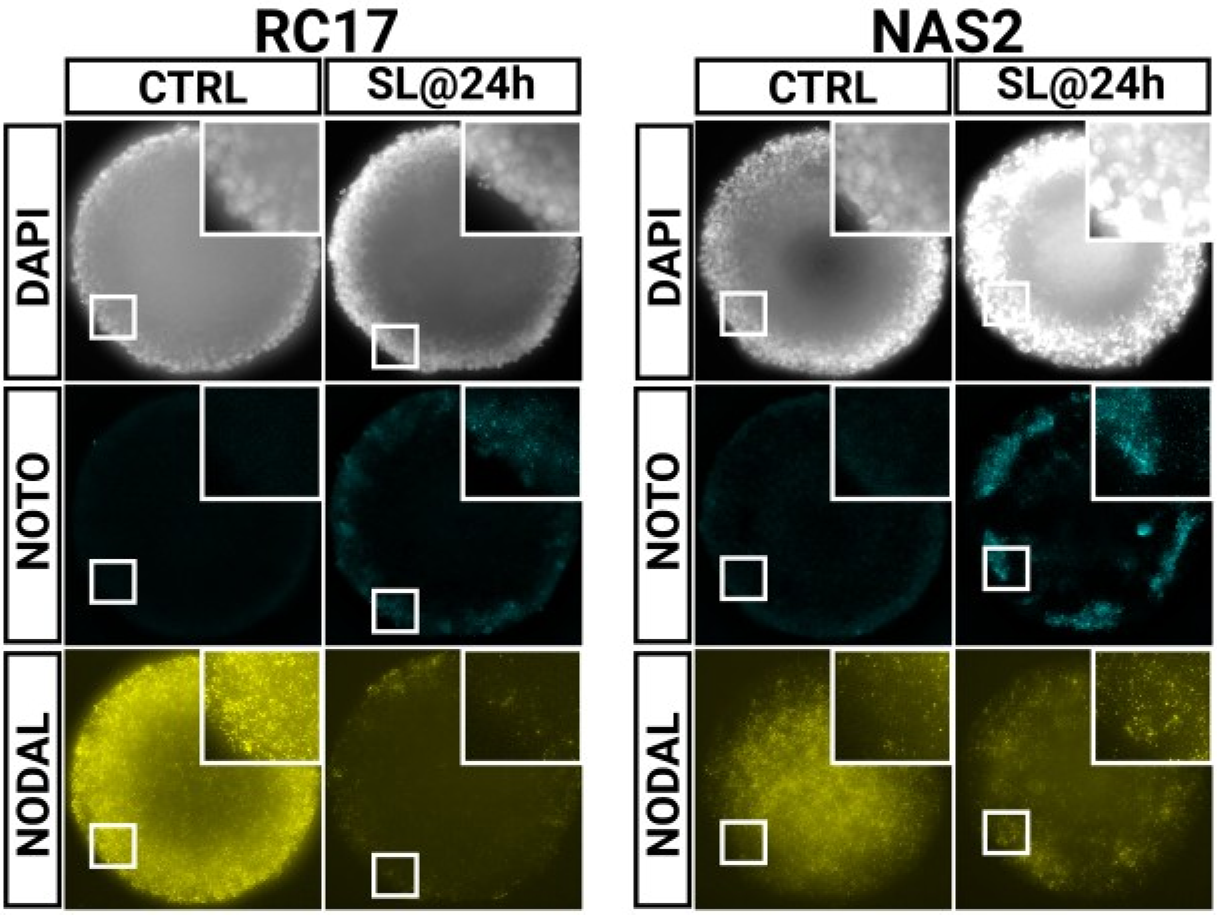


**Supplementary figure 5**

Representative widefield images of FISH staining against NOTO and NODAL transcripts in 500µm colonies at 48h of NotoPs induction (2µM CHIR, 20ng/ml FGF2 throughout and 10µM SB and 0.1µM LDN added at 24h). The data shows NOTO+ cells emerging at the periphery of colonies of 2 different cell lines.

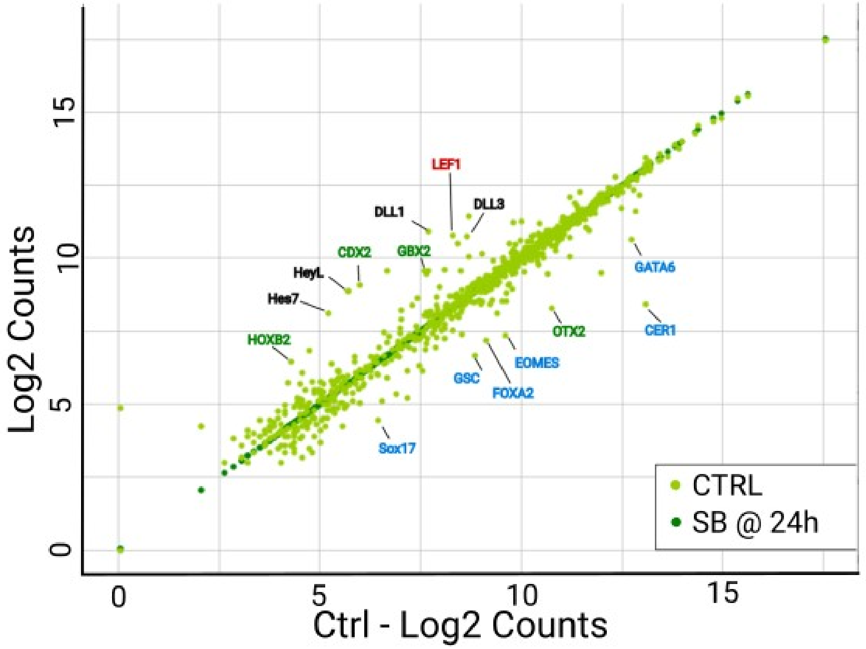


**Supplementary Figure 6**

Nanostring analysis of hESC grown on micropatterns in the presence of 2μM CHIR and 20ng/ml FGF2 for 2 days with or without 10μM SB added at 24h.

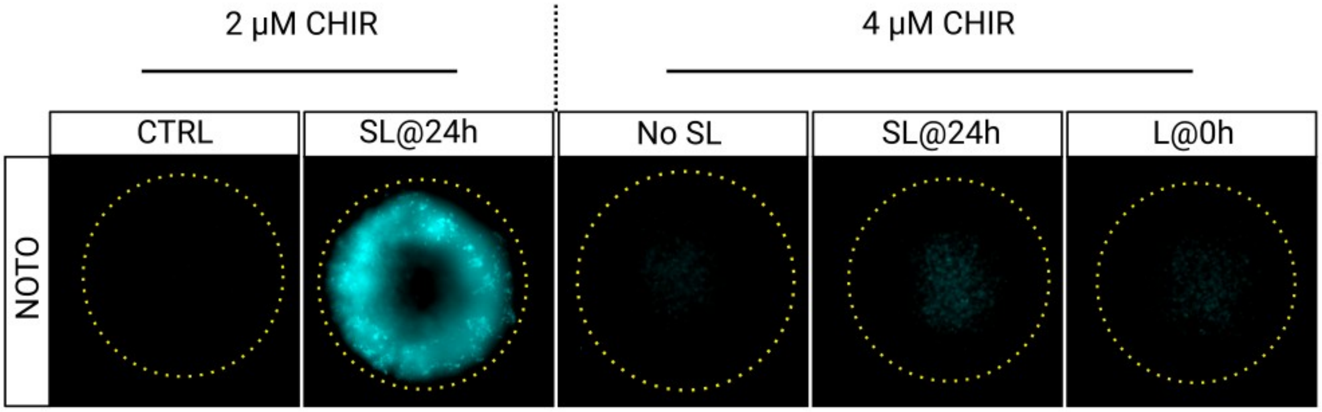


**Supplementary Figure 7**

Representative widefield images of FISH staining against NOTO transcripts in 500μm colonies at 48h of differentiation.
